# On the contribution of genetic heterogeneity to complex traits

**DOI:** 10.1101/2024.03.27.586967

**Authors:** Hai-Jun Liu, Kelly Swarts, Shuhua Xu, Jianbing Yan, Magnus Nordborg

## Abstract

Genetic heterogeneity, where different alleles or loci are responsible for similar phenotypes, reduces the power of genome-wide association studies and can cause misleading results. Although many striking examples have been identified, the general importance of genetic heterogeneity for complex traits is unclear. Here, we use a novel interpretative machine-learning approach to look for evidence of genetic heterogeneity in plants and humans. Our approach helps identify new loci/alleles influencing trait variation in several agriculturally important species, and we show that at least 6% of maize eQTL, half of them newly identified, exhibit evidence of allelic heterogeneity. Finally, we search for evidence of synthetic associations in human GWAS data, and find that as many as 3–5% may be affected. Our results highlight the need to take genetic heterogeneity seriously, and provide a simple approach for doing so.

---

The standard GWAS approach is a single-locus one—each variant is tested for association with the trait, and it is implicitly assumed that the presence of other causative loci does not affect marginal associations^1^. This is well-suited for identifying common variants with relatively large effect but is not designed for more complex situations^2,3^. Attempting to map multiple causal variants using single-locus models will generally decrease power and can bias estimates. In particular, it can generate so-called synthetic associations, where a non-causal variant ends up being the most significant by tagging the effects of multiple causal variants^4,5^. This phenomenon is augmented by close linkage, linkage disequilibrium, and epistatic interaction between causal loci, but does not require any one of them^5^. There are many beautiful examples in plants. For example, the sorghum *Sh1* gene that regulates seed-shattering^6^ was originally cloned through linkage mapping, but the confirmed causal mutations remained insignificant with standard GWAS, while several non-functional variants gained significance because of their strong linkage disequilibrium with a combination of a subset of causal mutations. Numerous other examples exist, also in non-domesticated plants like *Arabidopsis thaliana*^7–9^. Indeed it has been proposed that a general trend for top associations to be found some distance away from excellent *a priori* candidate genes exists due to synthetic associations^7,10^.

There is less evidence for synthetic associations in human genetics, but examples exist, such as the association between *NOD2* and Crohn’s disease^11^. A debate on the relevance of synthetic associations in human GWAS took place over a decade ago^4,12–14^, but it mainly focused on the role of rare variants. Whether synthetic associations matter in current, much larger GWAS^15^ remains untested, despite the growing evidence for multiple causal variants^16^.

In this study, we introduce a novel pipeline named GARFIELD (Genetic Aggregation with Random Forest and InterpretivE Logic Decisions), the basic idea behind which is illustrated in Fig. 1. Consider the simplest case of mutations at two different loci contributing to a phenotype, with either mutation being sufficient. At the population level, the effects of these variants are thus context-dependent and non-additive, and single-variant analysis may lack the power to detect either one. However, a pseudo-genotype representing the combined effect of the two loci would be significantly associated with the phenotype, and the “alleles” of this pseudo-genotype would reflect the underlying mechanism (Fig. 1a). This is the same logic that underlies the many “score” and “burden” tests that have been proposed to deal with large numbers of rare disease alleles, the difference being that we are mainly focusing on common alleles and make no assumptions about the direction of effect (nor do we utilize priors on which sites are likely to matter). An exhaustive recoding could be used to test for associations between the phenotype and the four pseudo-genotypes produced by any two variants to select the most promising one (Fig. 1b), but this is both computationally and statistically infeasible due to the vast number of tests, and we thus used a random forest algorithm^17^ to select variables exhibiting interactions (Fig. 1c). Simulations validated this strategy, demonstrating that our random forest implementation outperforms the standard linear mixed model in selecting real causal variants in simple models of genetic heterogeneity (Fig. S1). Finally, to make the results easier to interpret, GARFIELD uses logic gate terminology (e.g. SNP1 *or* SNP2; see Fig. 1d). When used on quantitative phenotypes, GARFIELD uses k-means clustering to identify two groups of phenotypes for generating binary pseudo-genotypes suitable for logic gate analysis. However, the raw quantitative trait values were still used during the association mapping step.

**Figure 1.**
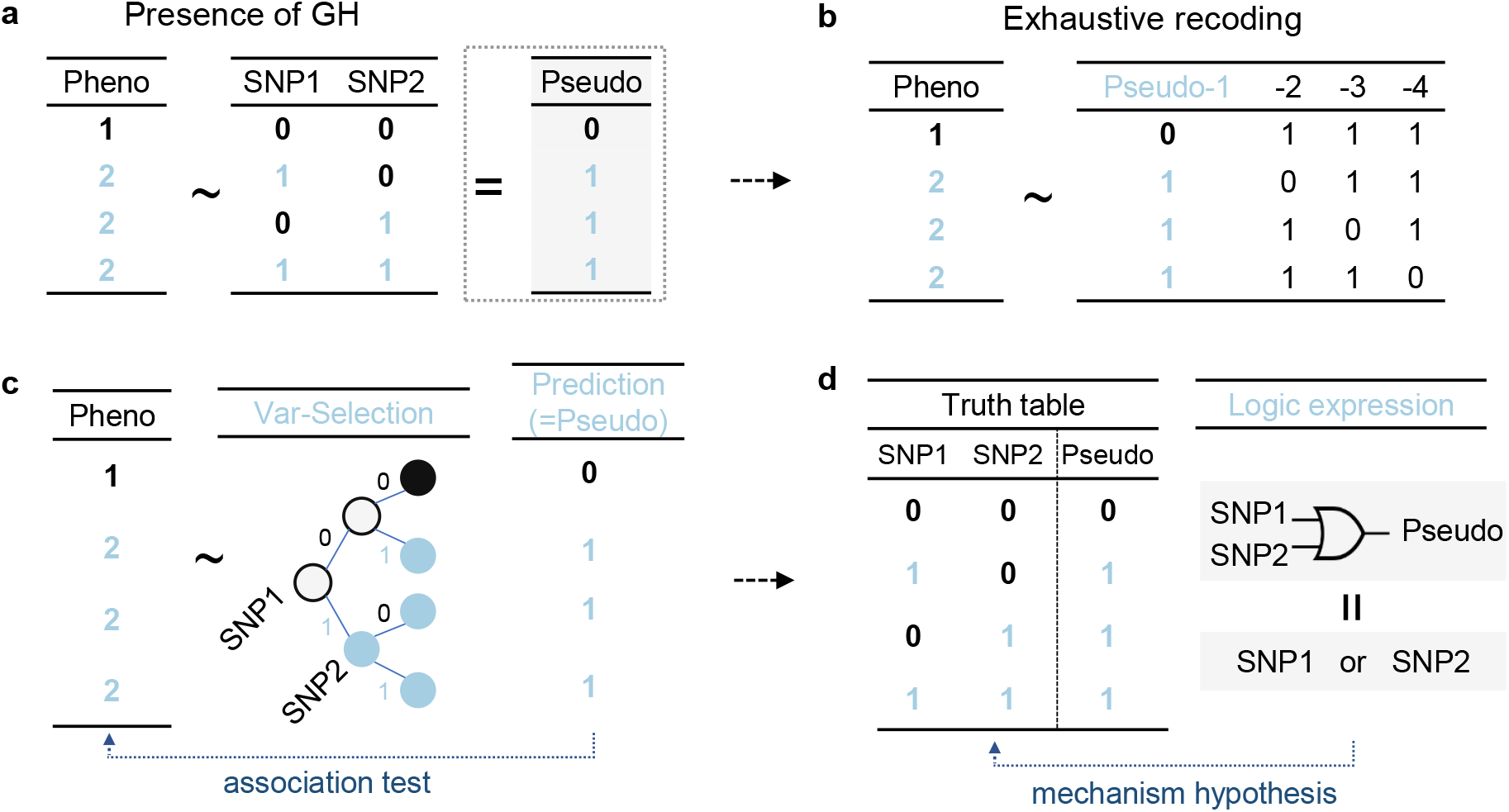
The rationale behind GARFIELD. **(a)** We illustrate our approach using the simplest case of a phenotype caused by a mutation at either of two causal loci in a haploid (or inbred diploid) species. The phenotype does not reflect the genotype at either locus, but is perfectly determined by a pseudo-genotype that combines both loci. **(b)** The “correct” pseudo-genotype could be identified by calculating all possible pseudo-genotypes, and choosing the most strongly associated one using GWAS. **(c)** A more efficient approach is to apply random forests to select variants with interaction effects. **(d)** Use of logic expressions to explain how the selected variants contribute to the pseudo-genotype.

In summary, GARFIELD combines the efficient variable selection (among thousands of markers from a given region or set of regions) and prediction advantages of random forests to produce pseudo-genotypes that help identify complex interactions, and subsequently use logic gates to explore and describe them. We note that several other methods for detecting allelic heterogeneity exist^3,18–20^, particularly various collapsing tests for capturing the cumulative effects of many rare variants^21,22^. Likewise, random forests have been used for variant selection for nearly 20 years^23,24^ (although our use of logic gates appears to be novel). In this paper, we introduce GARFIELD as a flexible tool that produces results that are easy to interpret, and our primary focus is not on methodology, but rather on exploring the evidence for heterogeneity in human and plant data. We use “genetic heterogeneity” in a very general sense, covering many different scenarios where a single-locus GWAS model might be inadequate.

We first demonstrate that looking for genetic heterogeneity in published data from several plant species increases power, identifies plausible interactions, and helps with fine-mapping; we then systematically look for evidence of synthetic associations in published human GWAS data, uncovering large numbers of plausible examples there as well.

## Results

### Allelic heterogeneity in agricultural traits

As a proof-of-principle, we re-analyzed the *Sh1* sorghum seed-shattering data that motivated this study—a beautiful example of allelic heterogeneity^6^. While single-variant GWAS failed to identify any of the known causal polymorphisms, GARFIELD identified two independent functional mutations, tagging all the non-shattering haplotypes (Fig. S2). Next, we carried out a genome-wide scan for allelic heterogeneity using kernel oil composition in over 500 natural lines of maize^25,26^. Single-variant GWAS had previously identified 26 loci; GARFIELD identified two more. Most convincing was *WRI1a*, a gene with a known role in regulating fatty acid synthesis^27,28^ and expression levels highly correlated with the trait. This gene had previously been assumed to be trans-regulated^25^, however, GARFIELD identified two *cis*-variants associated with both oil content and gene expression: a 2 kb indel in the 3’UTR and a missense SNP (Fig. 2). Note that the two variants are not in linkage disequilibrium, and are strongly associated with the phenotype without either variant being individually associated. In addition to *WRI1a*, GARFIELD revealed a very strong association with *GRMZM2G112149*, a methionine synthase2 gene, which is also a plausible candidate (Fig. S3).

**Figure 2.**
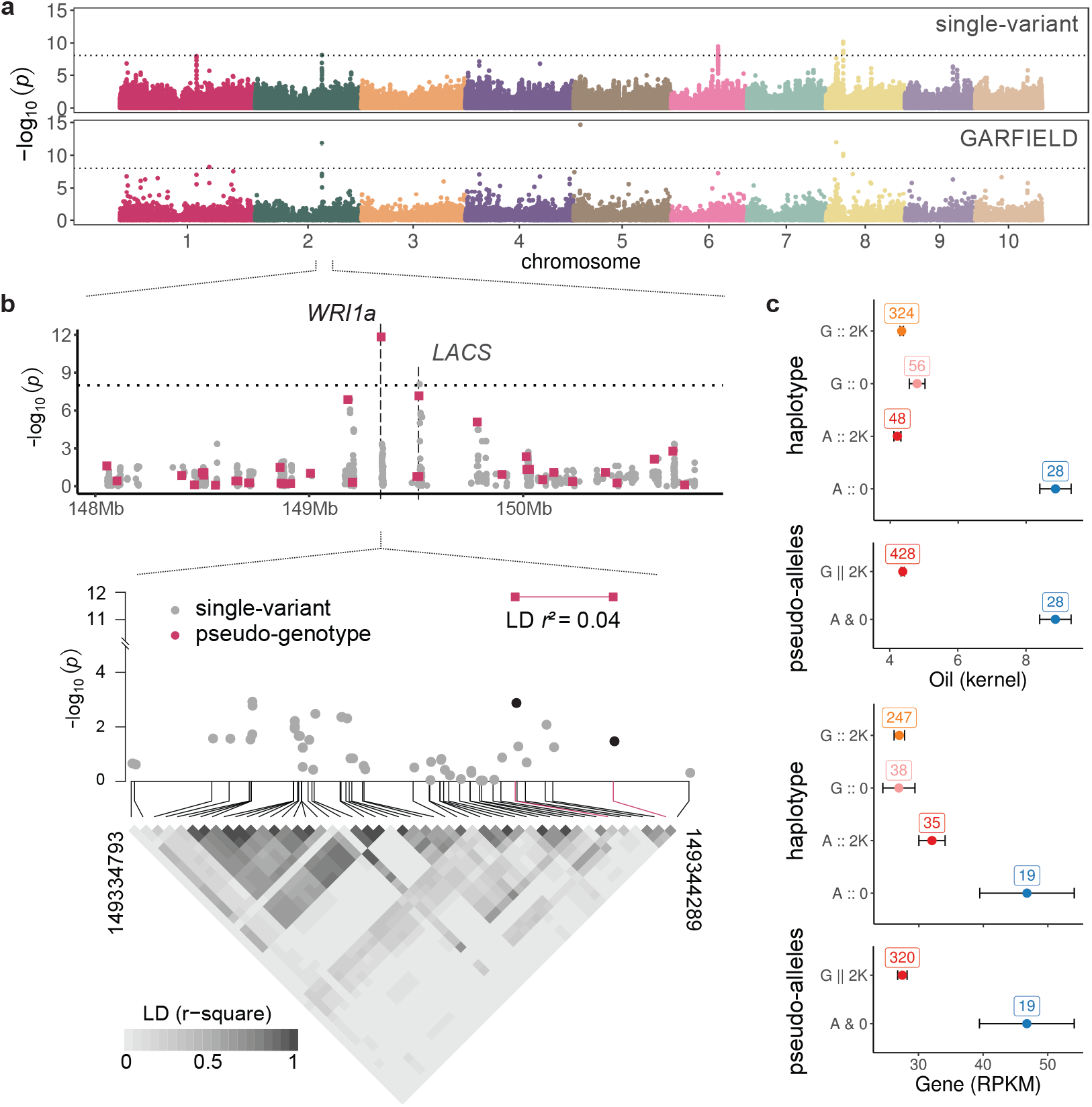
Allelic heterogeneity GWAS of maize kernel oil content. **(a)** GWAS of maize kernel oil content using single-variant GWAS and GARFIELD. The latter identified two new genes, *WRI1a* in chromosome 2 and *GRMZM2G112149* on chromosome 5. **(b)** Zoom-in on *WRI1a*, identified by combining a missense SNP and a 2kb indel at the 3’UTR, neither of which was marginally significant — while the two-locus combination is massively significant. **(c)** Mean oil content and gene expression across the four haplotypes and identified heterogeneity pseudo-genotype. Boxes show mean and standard error; the numbers in boxes give the number of individuals in each class.

Re-analysis of agronomic traits in rice also unveiled several associations not found with standard GWAS (**Table S1**). While some were supported by independent evidence, like *OsSPK* (Sucrose Synthase Protein Kinase; knock-down reduces storage products^29^) for grain weight (Fig. S4), and *Os-TAR1* (an IAA biosynthesis gene whose mutant phenotypes include reduced grain size and delayed filling^30^) for yield (Fig. S5), most are simply candidates requiring further validation.

Interestingly, even for loci already identified in standard analyses, heterogeneity analysis can potentially uncover new functional alleles. For instance, *OseIF4A2* (Os02g05330) emerged as most significant (*P* = 1.42 *×* 10^−11^ for SNP vg0202582923) in standard GWAS for grain weight, but GARFIELD revealed another SNP (vg0202588835, marginal *P* = 0.84) with minimal linkage disequilbrium (*r*^2^ = 0.01) that jointly with the previously identified SNPs was highly significant (*P* = 5.46 *×* 10^−13^; Fig. S6), explaining an additional 20% of effect size.

### Allelic heterogeneity in maize eQTL

To explore the general prevalence of allelic heterogeneity we need more phenotypes, and we therefore utilized gene expression data from seven tissues in a core maize diversity panel^31^. We identified 2,622 allelic heterogeneity eQTLs in 2,033 genes (6% of the total number of analyzed genes). This should be compared to a single-locus analysis, which identified 19,809 significant eQTL in 9,443 genes. Nearly half of allelic heterogeneity eQTL (1,219 in 1,038 genes) were found in genes without any significant association in single-variant analysis (**Table S2**). For genes with both single-variant and allelic heterogeneity associations, the top single-variant association was not included in a heterogeneity eQTL about onethird of the time (Fig. 3a), consistent with the former being a synthetic association (see Figs. S7-8 for examples). Using random combinations of expression and SNP data across genes, we estimate that our allelic heterogeneity approach had a false-discovery rate of less than 1%, lower than our estimate for single-variant analysis (Fig. 3b).

**Figure 3.**
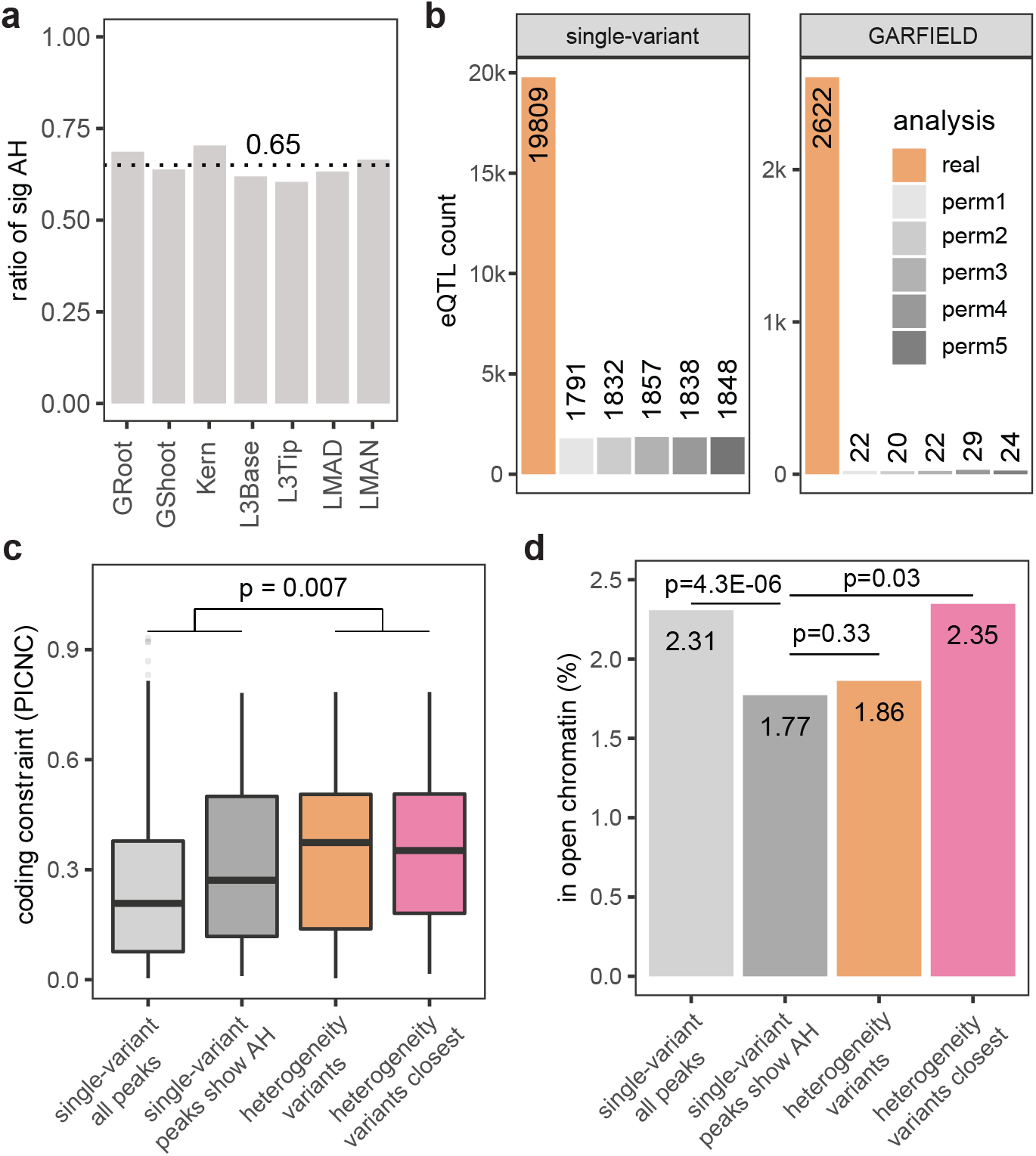
Using GARFIELD to reveal allelic heterogeneity in maize eQTL data. **(a)** Across seven maize tissues, for genes with both single-variant and allelic-heterogeneity eQTL, the top SNP identified by single-variant GWAS was not included in the multivariate associations roughly 35% of the times. **(b)** False-discovery rates for *cis*-eQTLs, estimatated by comparing eQTLs from five permutations of random genotype-phenotype gene pairs, *i*.*e*., the expression for a particular gene was combined with the SNP data (200 kb window) for a randomly chosen gene. **(c)** Evolutionary sequence conservation (PICNC scores) for variants identified using single-variant GWAS and using GARFIELD. For the latter, scores for all variants (heterogeneity variants) and for the one closest to the gene (closest) are shown. **(d)** Same as **(c)**, but gives the proportion of variants in open chromatin.

If a substantial fraction of single-variant eQTL were synthetic, we would expect the allelic heterogeneity eQTL to be more likely to be causal. We investigated this by considering the position of the identified SNPs with respect to annotation. In terms of coding constraint (PICNC) scores, allelic heterogeneity variants were significantly more constrained than variants identified using single-locus analysis (Fig. 3c). Only a small fraction of associated SNPs were in open chromatin^32^, and there was no general difference between variants identified using allelic heterogenity and single-locus analysis. However, single-locus variants in genes with evidence of allelic heterogeneity were less likely to be in open chromatin than single-locus variants in general (Fig. 3d).

### Genetic heterogeneity in pathways and among paralogs

Heterogeneity in the sense of this paper can also occur when mutations in different genes cause similar phenotypes, but systematically searching for such interactions is complicated by the vastness of the search space. However, it is feasible to carry out targeted searches, *e*.*g*., to search for members of a particular pathway. We tested this concept on *A. thaliana* flowering time data, specifically looking for interactions with the major flowering regulator *FLC*. For each gene in the genome, GARFIELD was run on a concatenated SNP set that covered the target gene region and *FLC*.

A total of 54 genes showed greater significance in combination with *FLC* than *FLC* itself (*P <* 5*×*10^−8^; **Table S3**). The second and third most significant associations were *GA2ox1* and *VIN3*, both well-known flowering-time regulators (Fig. S9). Overall, 12 of the 54 were known flowering genes from the FLOR-ID database^33^, a highly significant over-representation (*P* = 6 *×* 10^−61^). Ten of the 12 had previously not been discovered in single-variant GWAS^34^, and 4 were mapped to variants located within the gene (Fig. S9).

Another example of a targeted search for interaction is between paralogous loci, which frequently causes genetic redundancy^35–37^ — a type of interaction completely analogous to the simple model in Fig. 1 (the only difference is that it takes two or multiple mutations from distinct loci rather than one locus). We tested this on the rice data discussed above, applying GARFIELD to concatenated sets of closely related paralogous genes (see Methods; **Table S4**). This revealed several promising interactions. For grain weight, in addition to the newly identified allele at *OseIF4A2* (Fig. S6) discussed above, our gene-redundancy analysis revealed an interaction with its paralog *OseIF4A1*, which shows no marginal association (*P* = 0.44), but is highly significant jointly with *OseIF4A2* (*P* = 5.2 *×* 10^−10^; Fig. S6), explaining more of the trait variation (from 12.9% to 15.0%). Another example involves grain size, while *GW5* (Os05g0187500) has been cloned as the primary regulator for grain size^38–40^, our analysis revealed an interaction with its paralog *GW5L* (Os01g0190500) on grain thickness and width (Fig. S10), increasing the variance explained from 23.3% to 43.2%. This finding is consistent with knock-out and over-expression results indicating that *GW5* and *GW5L* function both independently and redundantly in regulating grain size and shape^41^. We looked for significant general enrichment of paralog vs non-paralog interactions but found none, consistent with genuine interactions being relatively rare (as well as, of course, a high fraction of false positives).

### Potential synthetic associations in human GWAS

Our results suggests that allelic heterogeneity is ubiquitous in plants. We would like to investigate humans, but this is complicated by the fact that individual genotype-phenotype data is generally not publicly available, especially for large systematic assessments. To get around this problem, we developed an approach to look for potential synthetic associations caused by allelic heterogeneity without having access to phenotypic data.

Briefly, we identified all variants within 100 kb of each published GWAS peak that were individually not strongly correlated with the peak SNP (*r*^2^ *<* 0.3, hence also not strongly associated with the phenotype used in the GWAS), and then used GARFIELD to search for combinations of these variants that *were* highly correlated with the peak SNP (*r*^2^≥0.8, hence potentially capable of explaining the phenotype; see Fig. 4a). We used this approach to mine the extensive collection of human GWAS peaks from the NHGRI-EBI GWAS Catalog^15^ and the deep variant profiles from the 1000 Genomes Project^42^ for associations that might be due to synthetic associations tagging multiple undetected causal variants.

**Figure 4.**
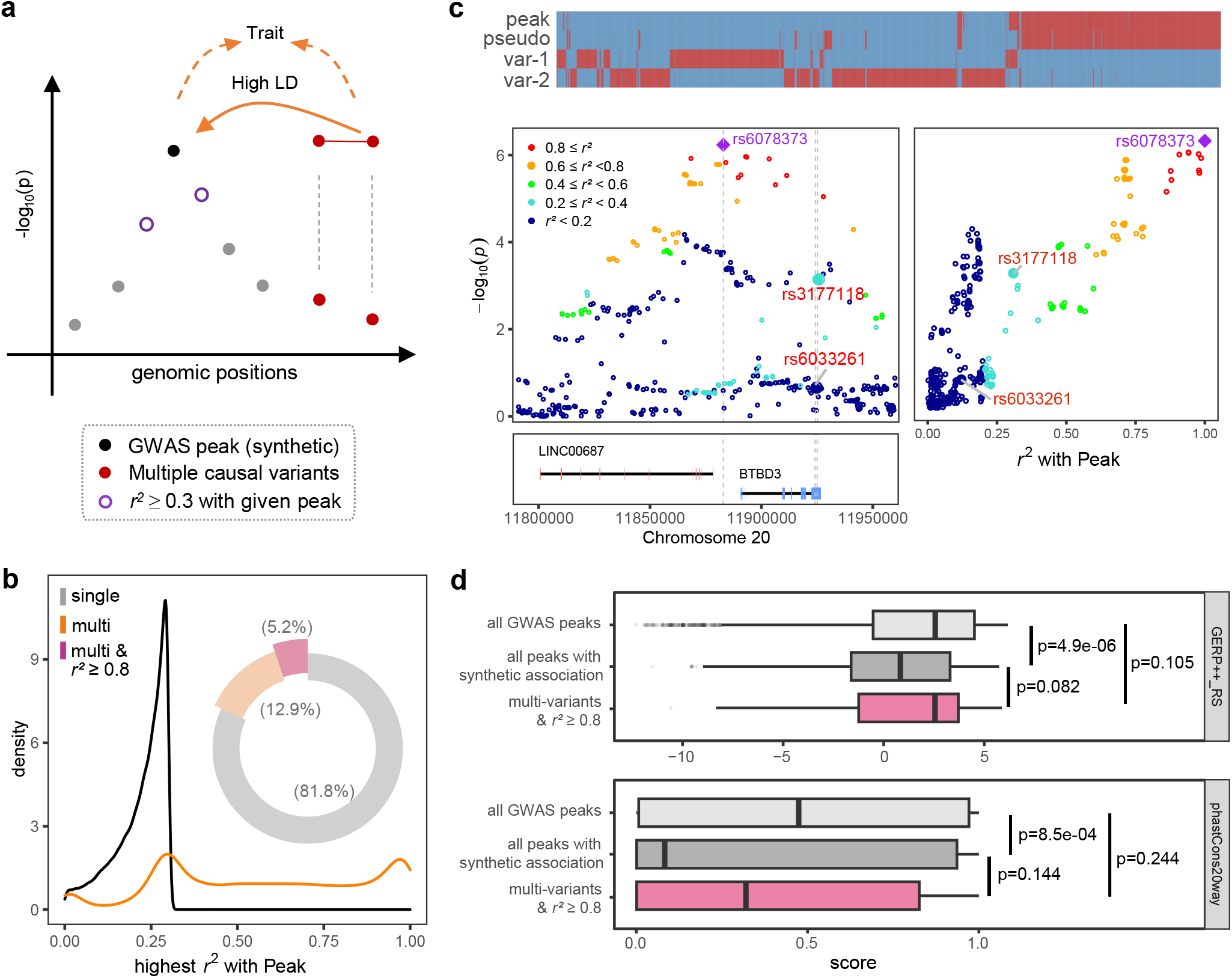
Possible synthetic associations in human GWAS. **(a)** To search for potential synthetic associations, we searched for SNPs within 100 kb of published associations that were individually not correlated with the peak SNP (*r*^2^ *<* 0.3), but which were jointly strongly correlated with it (*r*^2^≥0.8). **(b)** The orange curve shows the distribution of top multi-SNP *r*^2^ with the peak SNP across all 193,884 GWAS peaks in the “EUR” population (for other populations, see Fig. S12). The distribution has a bump at *r*^2^ ∼0.3, reflecting the fact that only SNPs with individual *r*^2^ *<* 0.3 were used. Also shown (black curve) is the density of single-SNP *r*^2^, by definition truncated at *r*^2^ = 0.3. For 81.8% of GWAS peaks, the best multi-SNP combination was less strongly correlated with the top SNP than the best single SNP (by definition *<* 0.3), but for 5.2%, the best multi-SNP combination was strongly correlated *r*^2^ *>* 0.8 (as shown in the pie chart). **(c)** An example of a potential synthetic association, involving a peak for smoking behavior in the EAS population. The three associated SNPs (only one of which is shown) are all strongly correlated with a combination of two SNPs inside *BTBD3*. The top panel shows the alleles at the peak SNP compared with the proposed pseudo-genotype, and its two constituent variants. Blue and red colors represent different alleles, and the x-axis shows haplotypes from all individuals. The left panel shows a regional Manhattan plot of the peak. The top SNP (rs6078373, in purple) is intergenic, but is strongly correlated with a combination of two SNPs (shown in red) in *BTBD3*. The right panel shows the marginal association of all SNPs with the phenotype as a function of their correlation with the top SNP. The y-axis is the same as the left. **(d)** Conservation scores for different groups of associated SNPs, including all analyzed GWAS peaks; all GWAS peaks with suspected synthetic association, and the heterogeneity variants that potentially cause synthetic association. Boxes show the 25%, median, and 75% of the distribution. Student’s t-test was used to evaluate significance.

Our procedure found no evidence of synthetic association in the great majority of GWAS results, however, depending on population, 3-5% matched our criteria (Fig. 4b). These numbers greatly exceeded those achieved when random variants (matched for allele frequency; Fig. S11a) were used as a control in the algorithm instead of the published GWAS hits (Fig. S12), suggesting that the pattern is unlikely simply to reflect background haplotype structure.

In total, our analysis identified 45,082 peaks as potential synthetic associations (**Table S5**) at least in one population analysis. Fig. 4c shows an example of an association with smoking and drinking behavior^43,44^. GWAS on these traits identified three intergenic SNPs around 11.88 Mb on chromosome 20. Our analysis showed that all three peaks are strongly associated with a pair of SNPs 40 kb away, within the coding region of *BTBD3*, a gene that promotes neural circuit formation^45^ and affects behaviors in mice^46^.

Another example of an intergenic peak that might be a synthetic association caused by variants in good candidate genes is 2:85315595, associated with high-density lipoprotein cholesterol levels, located between *TCF7L1* and *TGOLN2*, where heterogeneity analysis identified two variants within *TGOLN2* (one missense and one 3’UTR) that jointly are strongly correlated (*r*^2^ = 0.81) with the peak, suggesting that *TGOLN2* may be causal (Fig. S13). This seems plausible given that *TGOLN2* has been shown to significantly increase expression in response to EPA treatment^47^ and the related trans-Golgi network is crucial for the transport of high-density lipoprotein particles — while *TCF7L1* is expressed only in T-cells.

Two further examples of potentially synthetic associations that can be explained by pairs of variants in nearby genes are *TUT7* for cortical surface area (Fig. S14) and *ZKSCAN1* for height (Fig. S15). A very interesting example is the association between *APOE* and Alzheimer’s disease. While the importance of a major missense mutation (rs429358) is well established, another SNP located between *APOE* and *APOC1* was recently identified as a new independent (given *r*^2^ *<* 0.2 to rs429358) association with Alzheimer’s and over 40 other traits. However, we found that this SNP may be a synthetic association, as it is nearly perfectly correlated (*r*^2^ = 0.97) with two other SNPs: the above-mentioned missense mutation in *APOE* and one residing in a potential enhancer (E1957019) (Fig. S16). Further examples can be found in **Table S5**.

The examples above are consistent with the expected behavior of synthetic associations in that a single association is replaced by two (or more) that are more likely to be causal. To test whether this pattern is general, we considered evolutionary conservation using two different scores, GERP++ RS^48^ and phastCons20way mammalian^49^. For both scores, the associations we identified as potentially synthetic showed significantly less conservation than the full set of published associations—and also than the multiple SNPs we identified as potentially causing the synthetic association (Fig. 4d). This suggests that the published association was likely synthetic, and that our suggested replacements are more likely to be causal (although we note that some of these may of course also be synthetic). Further evidence that our findings are not simply random noise comes from the fact that evidence of synthetic association is very unevenly distributed across phenotypes (**Table S6**), as would be expected if there were true differences in genetic architecture, with some genes and pathways being more likely to harbor allelic heterogeneity.

Finally, we note that our findings do not support the notion that synthetic associations are generally common SNPs that tag multiple rare, independent disease-causing mutations^4^. The multi-variants we identified as potentially causing synthetic associations are generally more common than the corresponding peaks (Fig. S17). This suggests a completely different kind of epistatic interaction, where multiple mutations are required to cause the phenotype—an “AND” rather than an “OR” relationship—and the tag-SNP being one that ends up correlated with the multi-mutation haplotypes, necessarily rarer than the causal variants themselves.

## Discussion

In this paper, we use a simple but flexible algorithm based on random forests^17^ to search for combinations of genetic variants that jointly explain traits, but that are difficult to identify using standard, single-locus GWAS because of genetic heterogeneity. We show that many examples exist in plants, and argue that the phenomenon may be general in both plants and humans, with allelic heterogeneity affecting several percent of published associations. For these associations, the mapped variant is expected to be a “synthetic”, non-causal variant some distance away from the actual causal variants and not necessarily in linkage disequilibrium with any one of them^4,5^ — and we present evidence that this is indeed the case. Importantly, this is not just a statement that the top SNP is not causal, but that it is not even necessarily tagging a casual SNP. Furthermore, since heterogeneity is predicted to reduce power in addition to causing spurious associations, an unknown number of loci are likely to have been missed entirely. There is reason to take this problem seriously, especially when it can be done relatively simply, as this paper demonstrates.

We would argue that our findings are inherently plausible. While many decades of work in quantitative genetics have demonstrated that Fisher’s simple additive model works extremely well for breeding purposes and generally explains complex traits well^50^, this is expected given that effects are small and loci legion, and there is no reason to conclude from this that the model is generally correct at the level of individual loci — especially since there is abundant evidence of non-additive interactions from systems where effects are sufficiently large to allow accurate estimation. It follows from this that there is every reason to believe that a subset of GWAS results, obtained using a single-locus model that ignores interactions, are seriously misleading. Our results support this. Considering multi-variant models can help refine GWAS results and suggest more plausible causal variants. This is obviously important if the goal is to understand the molecular mechanism underlying a particular association, but it can also be important for understanding larger principles of genetic architecture. Two results are worth noting in this respect. First, since synthetic associations simply reflect random haplotype structure, they can be located anywhere. In both plants and animals, we find many examples where intergenic GWAS hits can be explained by multiple SNPs in genic regions. Indeed, this study was motivated by the first GWAS in plants showing a suspicious tendency for associations to be located near, but not near enough to, *a priori* candidates^7,10^. Uncritically treating GWAS hits as causal will introduce systemic bias against the regions of the genome that matter.

Second, our results suggest that synthetic associations in humans may often be caused by epistatic (synergistic) interactions between relatively common variants, rather than by multiple rare variants acting independently (in the molecular, not the statistical) sense. “AND” rather than “OR”, in other words. If this turns out to be true, then some diseases should be caused by rare haplotypes rather than rare alleles, and population genetic models of genetic architecture would have to consider multi-locus models of selection. Talking about the effect of individual alleles would no longer be sufficient — at least not in general.

Finally, we do not claim to have a general solution to this problem, nor do we propose that our results are unbiased.

Significant potential for algorithm improvement exists, including incorporating more powerful variable selection models (*e*.*g*., mixed random forests^51^), and broadening the search to genome-wide interactions. From a statistical point of view, GWAS poses an extremely challenging problem, but this is no excuse for not trying, and we hope to have demonstrated that much can be learned even with relatively simple methods.

## Methods

### GARFIELD

Random forests are computationally efficient and capable of capturing complex interaction effects^52^ but lack explicit interpretability. On the other hand, logic gates can provide clear relationships between variables^53^, but are relatively inefficient at analyzing a vast array of variables. GARFIELD integrates these two algorithms to leverage their respective strengths, using random forests for variant selection, and subsequently using logic gates to delineate probable relationships resulting in specific outcomes. Another innovation lies in employing the predictions/outcomes on selected combinations of variables, either from random forest or logic circuits, as pseudo-genotypes that represent intricate relationships. These pseudo-genotypes can be directly utilized in association mapping.

GARFIELD accepts binary phenotypes, or automatically conducts a k-means clustering to classify the best two groups for quantitative traits to enable subsequent pseudo-genotype generation and logic gate analysis. For genotypes there are specific modes, including gene-based, window-based, geneset-based, and ghost modes. In the gene-based mode, genotypes from the region of each gene, along with defined flanking regions, are considered. The window-based mode uses sliding windows with predetermined window and step sizes. For the gene-set-based mode, variants from a list of gene sets, along with extended intervals of each gene, are used. The genotypes in PLINK format were processed with [R genio v1.1.1](https://github.com/OchoaLab/genio).

In the current study, the gene- and window-based modes were employed to explore allelic heterogeneity. Paralog pairs were predominantly investigated using the gene-set mode, focusing on non-allelic heterogeneity in gene redundancy. This gene-set analysis could also encompass other types of gene groups, such as those based on pathways, protein-protein interactions, and more. The “ghost” mode was created specifically for investigating human synthetic associations, which closely resembled gene-based analysis, but with the variant itself treated as the “phenotype” and the other non-related variants (with correlation to top SNP lower than a specified threshold) were used as input genotypes within a defined window.

The random forest analysis (implemented using the “ranger” library^52^, v0.14.1) was initially employed to estimate the importance of variants (with “minnode = 5, seed = 222”) for the given phenotype. This analysis was carried out for each predefined variant list, determined by the different modes described above. A custom function was applied to select the top variables, focusing on the points of the greatest absolute difference in importance scores. In each analysis (*e*.*g*., one gene or one window), a maximum of 5 variants was selected. This limit was used because we considered more than 5 causal variants unlikely, and in any case almost certainly undetectable given the available sample sizes.

When a single best variable was selected (typically the peak variant identified through conventional single variant-based analysis), its original genotype was directly adopted for further use. In the scenario where multiple variants were chosen, these selected variants were utilized to retrain the prediction model within the random forest framework. The resulting predictions from this model were then used as a new variable termed the “pseudo-genotype.” This pseudo-genotype encapsulated the most significant associations among the chosen multiple variants and the phenotype.

To establish an interpretative link between the chosen multiple variants and the pseudo-genotype, another crucial step was the derivation of the most optimal Boolean logic expression. This expression aimed to effectively capture the connection between the genotypes of the selected multiple variants and the pseudo-genotype as output. This inference process was achieved through logic regression, which was executed using the logicFS package^53^ (v2.2.0). Specifically, the logic.bagging function was employed with parameter settings “B = 100, nleaves = 10, ntrees = 1, seed = 222, start = 2, end = −2, iter = 1000”.

In this analysis, the disjunctive normal form (DNF, for example, var A or NOT var B and var C) of a logical formula was utilized to represent the logic expression. This form was chosen due to its suitability for describing the likely interpretative relationships. Note that while the selected variants and pseudo-genotypes predicted by the random forest method might not always be entirely explainable through logic expressions alone, another prediction was employed to update the pseudo-genotypes based on the determined logic expression, akin to the approach taken for the random forest above. This process aimed to guarantee that the logic expressions could capture the outcomes arising from the interaction of the selected multiple variants with their corresponding pseudogenotypes.

In the end, GARFIELD outputs the DNF that describes the relationship between the selected variants and their corresponding pseudo-genotype. These pseudo-genotypes can then be utilized to test their association with the provided phenotypes (the raw continuous values should be used in the case of quantitative traits) like any other genotype, with the DNF providing insight into possible underlying mechanisms. In cases where only one variable was selected, the DNF would correspond to the individual marker itself, and the pseudo-genotype would remain identical to its original raw genotype. An option can be set in GARFIELD to filter out these instances as they provide fully redundant information compared to conventional single marker-based analysis. However, in the present study, all these cases were retained for further analysis.

### GWAS

Associations between the pseudo-genotypes output by GARFIELD and the phenotypes can be investigated using standard association mapping methods. In this study, both single marker and genetic heterogeneity-based mapping were conducted using EMMAX^54^. For these analyses, the top 10 principal components (estimated using PLINK^55^) for plants together with the IBS kinship matrix (obtained from emmax-kin with parameters “-s -d 3 -S 222”) were incorporated. When relevant, the pseudo-genotypes were added to the original genotypes to calculate the PCs and kinship matrix.

To establish significance thresholds, the effective variant number (*n*_eff_) was estimated using the Genetic Type 1 Error Calculator^56^, and the p-value thresholds were adjusted to 0.01/*n*_eff_. This led to thresholds ranging from 9.2*×* 10^−9^ to 2.4*×*10^−8^ depending on species (maize, rice, and *A. thaliana*) and phenotype (sample sizes varied slightly). For simplicity, a threshold of 1 *×* 10^−8^ was adopted for all traits across all species.

The same threshold was used for the heterogeneity analyses. The number of additional markers added to these analyses was trivial (typically fewer than 20k), and all simulation and permutation analyses showed a very similar distribution of p-values to that of single-marker analysis, with no indication of a higher false-positive rate. That said, all cases mentioned in the main text that were exclusively identified in the heterogeneity analysis exhibited significance levels several orders of magnitude higher than those in the single-variant analysis.

### Simulating multiple causal variants

We used simulations to compare the performance of GARFIELD and single-variant GWAS in the presence of multiple causal variants. The “–simulate” command in PLINK was used to generate random samples of 400 individuals, with 50 variants per locus, and 1 or 2 causal variants within each locus. Linkage disequilibrium between the two causal variants was set to be “independent” (*r*^2^≈ 0).

Ten independent genotype sets were generated for the simulation, varying the frequencies of causal polymorphisms from 0.02 to 0.3 with a step of 0.02. Furthermore, the “–simu-qt” in GCTA^57^ (v1.93.0) was employed to simulate case-control phenotypes based on the generated genotypes, incorporating the designated causal variants with a specified heritability of 40% and disease prevalence of 50%. For each genotype set, a total of 50 simulated phenotypes were generated. The kinship matrix was utilized in association mapping to assess the efficacy of each scenario, encompassing both single variant and random forest analysis. No PCs are considered as covariates here, since no population structure is simulated. All other analyses and thresholds were performed for the real data. The power of the test was evaluated by determining the fraction (across a total of 50 repeated phenotypes) of accurately identified causal variants. The simulation results were in line with earlier discoveries that highlighted the efficacy of random forest in identifying variants exhibiting both additive effects and interactions^58,59^.

### Maize oil content

We used a population of 540 individuals genotyped for 1.25M (minor allele frequency, MAF≥5%) markers, using a combination of RNA-seq, 50K array, 600K array, and GBS data^26^. The genotypes are available from [Maizego](http://www.maizego.org/Resources.html) or the Modem database^60^. The total oil content of the same population^25^ was studied, and the re-sequencing data for the *WRI1a* locus was obtained from the authors. The corresponding B73 v2 annotation was used for this genotype. Paralog information was retrieved from the BioMart tool of [Ensemble Plants](https://plants.ensembl.org/index.html) and paralog pairs showing an identity greater than 0.7 in both directions (query to target and target to query) were retained.

Variants with a missing rate exceeding 20% were filtered out, and the remaining missing calls were imputed using Beagle^61^ (v5.2). Variants with an MAF below 2% (after considering the overlap of variable line numbers in different traits) were excluded from further analysis. A flanking region of 10 kb for each gene was used for gene and paralog analyses. A window size of 100 kb with a step size of 20 kb was used for window-based analysis.

The *WRI1a* locus was independently analyzed using both gene and window-based approaches because the number of individuals re-sequenced differed from the entire genotype panel that shared overlaps with oil traits (456 vs. 476). The association results were subsequently integrated into the whole genome analysis.

### Rice agronomic traits

Phenotypes and genotypes were retrieved from [Rice-VarMap v2.0](http://ricevarmap.ncpgr.cn/download/)^62^. A total of 529 accessions were examined for 10 agronomic traits: Heading date, Plant height, Num panicles, Num effective panicles, Yield, Grain weight, Spikelet length, Grain length, Grain width, and Grain thickness. The analyses involved using both MSU release 7 and IRGSP annotations for gene- and paralog-based investigations, respectively. Rice paralog pairs were acquired from [Ensemble Plants BioMart](https://plants.ensembl.org/index.html), with retention of pairs exhibiting identity greater than 0.7 in both query-to-target and target-to-query directions.

Lines missing over 50% of genotypes from these 529 lines were excluded. The remaining missing genotypes were imputed using Beagle 5.2, and only genotypes with an MAF of 5% were selected for further analysis. For gene and paralog analyses, a flanking region of 10 kb for each gene was incorporated. In window-based analysis, a window size of 50 kb with a step of 20 kb was used.

#### *A. thaliana* flowering time

SNPs derived from whole-genome sequencing of 1,135 lines and flowering time in 10^*°*^C growth chambers (FT10mean)^34^ were utilized. The genotypes were downloaded from 1001 Genomes. Missing SNPs were imputed using Beagle 5.2, with retention of SNPs featuring an MAF of 2% for subsequent analyses. All genes annotated in [TAIR10](https://www.arabidopsis.org/) were paired with *FLC* for non-allelic heterogeneity investigations, with a flanking region of 20 kb included for each gene. A significance threshold of *P <* 5*×* 10^−8^ was used for this analysis. Enrichment for known flowering regulators was calculated using the 306 genes listed in the Flowering Interactive Database, [FLOR-ID](http://www.phytosystems.ulg.ac.be/florid/)^33^.

### Maize multi-tissue transcriptome data

Our search for evidence of allelic heterogeneity in maize eQTL results was based on 300 individuals that had undergone RNA-sequencing across seven tissues. The genotype and expression data are publicly available at [Data Commons](https://datacommons.cyverse.org/browse/iplant/home/shared/commons_repo/curated/Kremling_Nature3RNASeq282_March2018), and include the imputed genotypes from Maize Hapmap v3.2.1 and the expression matrix `TASSEL_fmt_expression_w_covariates`. The corresponding B73 v3 annotation was used.

The tissue codes are: GShoot, germinating shoot; GRoot, germinating root; L3Base, base (2cm) of the third leaf; L3Tip, tip (2 cm) of the same third leaf; Kern, kernels from the center of cobs; LMAD, Leaf Mature Adult Day; and LMAN, Leaf Mature Adult Night. For more details, please see the Readme filed or the original manuscript^31^.

For each gene within each tissue, a 100 kb flanking region was included to conduct allelic heterogeneity analysis, which was carried out as described above except for the incorporation of 25 PEER hidden factors as covariates^31^.

To assess the significance of signals of allelic heterogeneity affecting *cis*-regulation of expression, we used permutation. We randomly shuffled the genotype-phenotype combinations so that, for each gene’s expression, genotypes from other, randomly chosen genes within the same 100 kb region were used in the eQTL mapping. This random pairing sampled without replacement, so that each gene was paired with another gene only once. False positive rates were then estimated by comparing the number of significant eQTLs in the permuted and the original data.

To compare the functional potential of variants identified using GARFIELD and single-locus methods, two different approaches were used. First, to cover the noncoding genome, we considered the presence in open chromatin, specifically MNase-hypersensitive regions obtained from shoot tissue were^32^. Second, for coding regions, we employed the PICNC (Prediction of mutation Impact by Calibrated Nucleotide Conservation)^63^. A higher PICNC score indicates a stronger degree of evolutionary constraint, and these metrics of the maize genome are available from [Data Commons](https://datacommons.cyverse.org/browse/iplant/home/shared/commons_repo/curated/Ramstein_SNPConstraintPrediction_2022).

### Potential synthetic associations in human GWAS peaks

Our study was based on published human GWAS results from the [NHGRI-EBI Catalog](https://www.ebi.ac.uk/gwas/) (data released on 2023-07-05)^15^. All peaks with demonstrated SNP-SNP interactions (multiple variants separated by “x”) and multi-SNP haplotypes (multiple variants separated by “;”) were removed. Peaks not represented in the genotype data of the 1000 Genomes Project (spanning chromosomes 1 to 22 only), or exhibiting multiple alleles, or having an MAF lower than 2%, were also eliminated. This left a total of 206,609 unique variants and 382,504 genotype-phenotype associations (variant*×*PUBMEDID *×* DISEASE/TRAIT). The summary statistics were also downloaded from the GWAS Catalog, with the exception for association statistics for human height which were downloaded from the [GIANT Consortium](https://www.joelhirschhornlab.org/giant-consortium-results) (all population groups were used).

The phased genotype data from the high-coverage wholegenome sequencing of the 1000 Genomes Project^42^ was obtained from [EBI](http://ftp.1000genomes.ebi.ac.uk/vol1/ftp/data_collections/1000G_2504_high_coverage/working/20220422_3202_phased_SNV_INDEL_SV/) (downloaded at 14/7/2023). For further analysis, the dataset was refined to include only biallelic variants located on chromosomes 1 to 22, with a MAF of at least 2%. This genotype dataset has no missing calls, thus eliminating the need for imputation.

Given that phased information is often disregarded during linkage disequilibrium analysis, and for the sake of consistency with the binary nature of logic gates analysis, diploid phased genotypes (0|0, 0|1, 1|0 and 1|1) were transformed into two haploid individuals. This approach resulted in each variant only having two possible values (0 or 1). The “phenotype” (the GWAS peak genotypes) was processed in the same way. Note that GARFIELD-generated pseudo-genotypes reflecting potential synthetic associations, as well as all haploid genotypes, can be easily combined back into the original diploid phased genotypes. The linkage disequilibrium estimates obtained from haploid genotypes displayed a high degree of correlation with those being merged back to diploid genotypes (Fig. S11b), demonstrating the robustness of findings from haploid analysis.

For each GWAS peak, the analysis all variants within a 100 kb flanking region that were not in strong linkage disequilibrium with the peak (*r*^2^ *<* 0.3). We further pruned the data to eliminate any variants displaying strong linkage disequilibrium (*r*^2^≥0.9) with another. This step was aimed at enhancing the power of identification of the important variants during random forest analysis. The presence of strong linkage disequilibrium between variants would reduce the likelihood of both variants being selected, as they would share and consequently lower the importance score.

Considering the genotype of each peak variant as a “phenotype”, we used GARFIELD to screen all other filtered variants association, distinguishing between three different outcomes based on the nature of the strongest one:

1. A single variant, for which *r*^2^ *<* 0.3 with the peak (by necessity as more strongly associated variants had been removed).

2. Multi-variant, moderately associated (*r*^2^ ≤ 0.8).

3. Multi-variant, strongly associated (*r*^2^ *>* 0.8).

The analysis was run on the full sample of 2,504 individuals (referred to as the “human” group), and within each of the five subsets, specifically African (AFR), Admixed American (AMR), East Asian (EAS), European (EUR), and South Asian (SAS). And those cases identified by a mix of single variant and Multi-variants, that either the “single” or “multi” show indistinguishable association levels to the peaks, were excluded in further analysis.

To estimate the rate of spurious associations produced by our analysis, GARFIELD was also turned using randomly selected control variants as “phenotype” instead of the actual GWAS peak. These control variants were identified using PLINK with parameters “–maf 0.02 –indep-pairwise 500 50 0.3”, and a second iteration was performed employing “–indep-pairwise 500 500 0.3”. From these pruned variants, control variants were then randomly chosen to match the actual GWAS peaks in MAF and number (illustrated in Fig. S11a).

Upon careful examination of the results, we identified instances where the same haplotype combination from multiple selected variants yielded different alleles of the pseudogenotypes (refer to Fig. S11c). This occurrence might have arisen due to other hidden variants or unaccounted factors. To address this, a filtering process to exclude these situations was implemented for both GWAS peaks and control variant studies.

For intricate DNF expressions (such as snp_A & snp_B & snp_C|| snp_A & snp_B & snp_D, where snp_D typically exhibits high linkage disequilibrium with snp_C, and snp_A & snp_B & snp_C displays fairly similar association to the target “phenotype”), a simplified expression (in this case, snp_A & snp_B & snp_C) was adopted for further analysis, to explore the frequency distribution and the LD between each variant comprising heterogeneity and the peak genotypes (Fig. S17). To compare the performance of the methods in terms of identifying likely causal variants, predicted variants effects were downloaded from [EBI](http://ftp.1000genomes.ebi.ac.uk/vol1/ftp/data_collections/1000G_2504_high_coverage/working/20201028_3202_raw_GT_with_annot/). We used two quantitative metrics that measure conservation (a higher score indicates a greater degree of conservation):

1. phastCons20way_mammalian represents the phastCons conservation score (range from 0 to 1) based on multiple alignments of 20 mammalian genomes including humans;

2. GERP++_RS score (range from −12.3 to 6.17 in this dataset) assesses evolutionary conservation by evaluating the pattern nucleotide substitutions over evolutionary epochs.

It is worth noting that we employed a pruning approach for highly linked genotypes during the search for heterogeneity for each GWAS peak. While this would enhance the power to identify potential heterogeneity, it will potentially interfere with variant effect analysis here since highly linked variants were chosen randomly.

Finally, the MAPPED TRAIT URI column in the GWAS Catalog was utilized to identify trait/disease ontologies, encompassing Experimental Factor Ontology, Human Phenotype Ontology, and Mondo Disease Ontology. Entries lacking ontology information were excluded. The [Ontology Lookup](https://www.ebi.ac.uk/ols/index) Service was explored for comprehensive information about each ontology term. To identify phenotypes enriched in potential synthetic associations, trait ontology terms of peaks identified as potential synthetic associations were compared against terms from all the studied peaks. The Hypergeometric test (implemented through the “phyper” function in R with “lower.tail = FALSE”) was employed, followed by multiple comparison corrections, and terms with a false discovery rate (FDR) of 5% or lower were deemed significantly enriched. Ontology terms including at least two potential synthetics associations were retained.

## Data and code availability

All the genotypes and phenotypes utilized in the analysis were obtained from publicly accessible sources, as outlined in the respective methodology sections. GARFIELD is freely available on [GitHub](https://github.com/heroalone/Garfield).

## Acknowledgements

We would like to thank Doc Edge, Jianming Yu, Wenyu Yang, Tom Ellis, and Almudena Mollá Morales for their helpful comments and feedback on the manuscript. This project has received funding from the European Union’s Framework Programme for Research and Innovation Horizon 2020 (2014-2020) under the Marie Curie Skłodowska Grant Agreement Nr. 847548 (to H-J.L.). S.X. also acknowledges the support from the National Key Research and Development Program of China (2023YFC2605400) and the National Natural Science Foundation of China (NSFC) grants (32030020).

## Author contributions

H-J.L., M.N., and J.Y. conceived and designed the analysis. H-J.L. performed data collection and analysis. H-J.L., M.N., J.Y., K.S., and S.Xu. interpreted the results. H-J.L. and M.N. drafted the manuscript, with revisions and approval by all authors.

## Competing interests

The authors declare no competing interests.

## Supplementary information

### This file includes Supplementary Figures 1-17

**Supplemental Figure 1**. The performance of GARFIELD compared to single-variant mixed model in simulations.

**Supplemental Figure 2**. Application of GARFIELD to sorghum *Sh1* seed-shattering data.

**Supplemental Figure 3**. Allelic heterogeneity GWAS of maize kernel oil content.

**Supplemental Figure 4**. Allelic heterogeneity GWAS of rice grain weight identifies *OsSPK*.

**Supplemental Figure 5**. Allelic heterogeneity GWAS of rice yield identifies *OsTAR1*.

**Supplemental Figure 6**. Allelic heterogeneity GWAS of rice grain weight identifies a novel allele of *OseIF4A2* locus and an interaction with its paralog, *OseIF4A1*.

**Supplemental Figure 7**. Examples of allelic heterogeneity eQTL in maize data.

**Supplemental Figure 8**. Instances of heterogeneity eQTLs for the same gene in different tissues.

**Supplemental Figure 9**. Identification of flowering time genes by non-allelic heterogeneity analysis on gene pairing with *FLC*.

**Supplemental Figure 10**. Identification of rice *GW5* paralogs associated with grain shape.

**Supplemental Figure 11**. Technical issues in the analysis of human synthetic associations.

**Supplemental Figure 12**. Identifying potential synthetic associations in human GWAS.

**Supplemental Figure 13**. Potential SA in human GWAS at *TGOLN2* locus.

**Supplemental Figure 14**. Potential SA at *TUT7* locus.

**Supplemental Figure 15**. Potential SA of *ZKSCAN1* on body height.

**Supplemental Figure 16**. Potential misleading findings discovered at *APOE* locus to Alzheimer’s disease.

**Supplemental Figure 17**. Additional features of potential synthetic associations in human GWAS.

**Another independent file “Supplemental Tables**.**xlsx” for supplementary Tables 1-6:**

**Supplemental Table 1**. List of allelic heterogeneity loci of 10 agronomic traits in rice.

**Supplemental Table 2**. List of allelic heterogeneity eQTLs in maize.

**Supplemental Table 3**. Detection of *FLC* partners in FT10.

**Supplemental Table 4**. List of non-allelic heterogeneity associations between paralogs in rice.

**Supplemental Table 5**. List of potential SA in human GWAS.

**Supplemental Table 6**. Phenotype Ontology (PO) enrichment of human potential SAs.

**Supplemental Figure 1.**
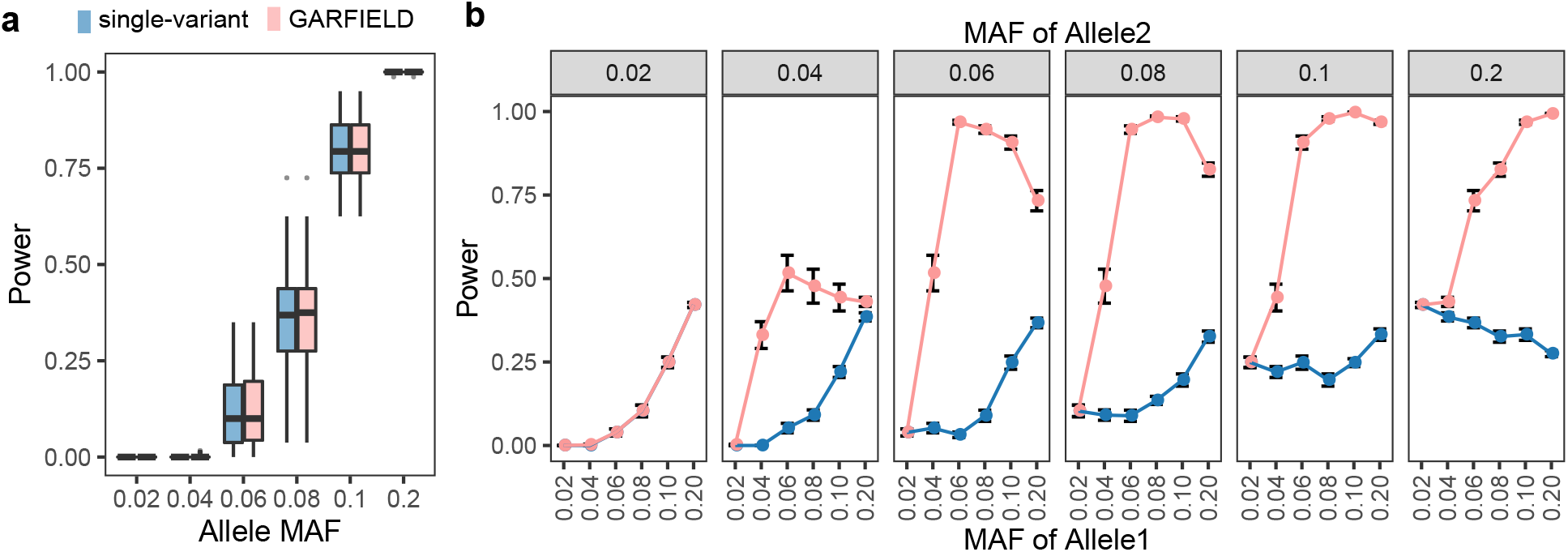
The performance of GARFIELD compared to the single-variant mixed model in simulations. **(a)** In simulations (see Methods) with a single causal variant, GARFIELD had similar power to detect the causal variant as the standard single-variant mixed model. The bar in the boxplot shows the median and the box represents the first to the third quartile. **(b)** In simulations of allelic heterogeneity with two causal variants. Power here is defined as the fraction of variants correctly identified. When one variant is rare, the simulation effectively reduces to a single-variant model, and both methods perform equally well, as in panel **a**. In all other cases, GARFIELD outperforms the single-variant model, which rarely identifies more than 1 of the variants (and is hence limited to 50% power). The dots and error bars show the mean and standard error, respectively.

**Supplemental Figure 2.**
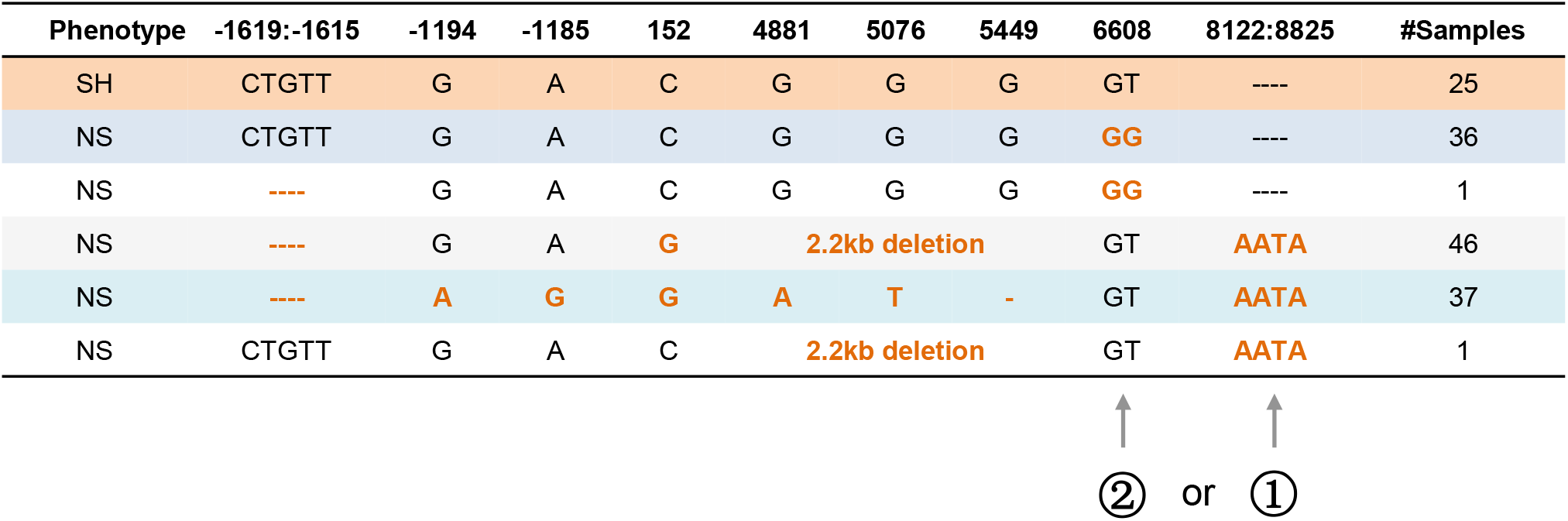
Application of GARFIELD to sorghum *Sh1* seed-shattering data. Non-shattering evolved multiple times during sorghum domestication^6^. Mutant alleles known to be involved are highlighted in orange. GARFIELD successfully identified two of these alleles, which together fully distinguished shattering from non-shattering phenotypes. Specifically, the insertion at position 8122-8825 and the SNP at 6608 affecting splicing, emerged as the first and second most important variants in the random forest analysis, respectively, mainly due to the higher frequency of the insertion than the splicing SNP.

**Supplemental Figure 3.**
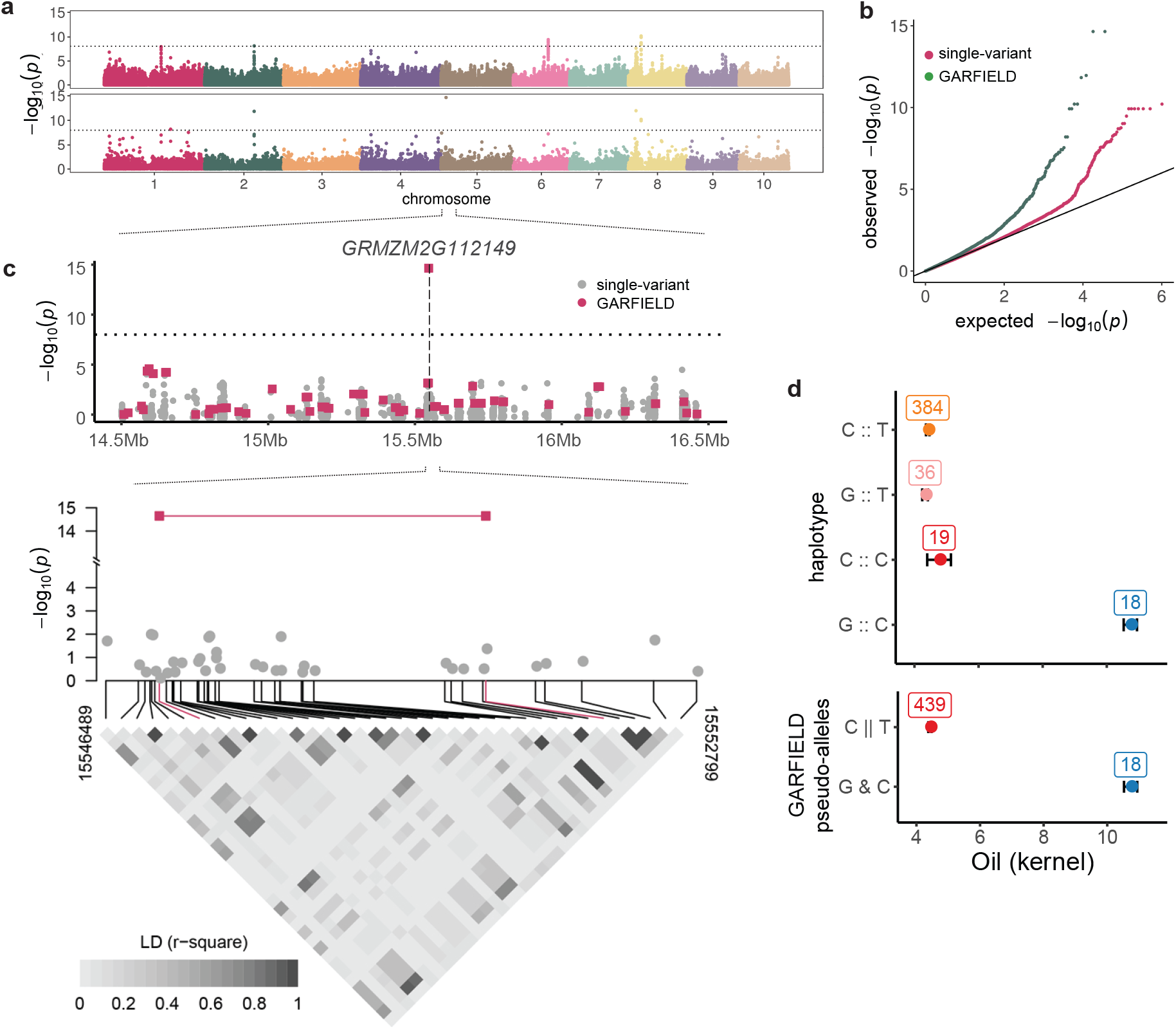
Allelic heterogeneity GWAS of maize kernel oil content. **(a)** GWAS of maize kernel oil content using single-variant GWAS and GARFIELD (same as in Fig. 2). **(b)** Quantile-quantile plot of p-values from both methods. **(c)** Zoom-in on *GRMZM2G112149*, a methionine synthase2 locus exclusively detected by GARFIELD. **(d)** Mean oil content across the four haplotypes and identified pseudo-genotype.

**Supplemental Figure 4.**
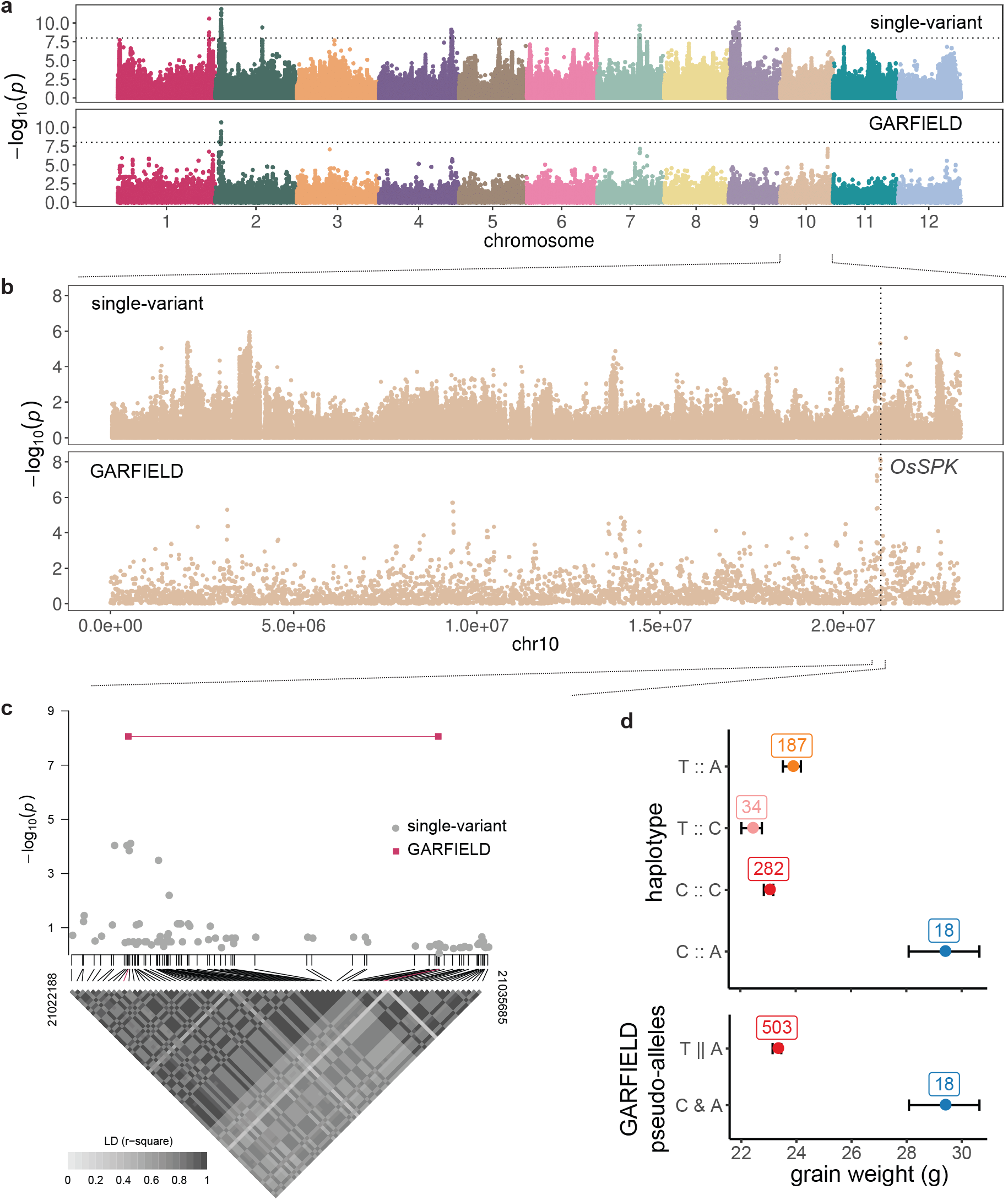
Allelic heterogeneity GWAS of rice grain weight identifies *OsSPK*. **(a)** Comparison of single-locus GWAS and GARFIELD. **(b-c)** Zoom-in on *OsSPK*. **(d)** The grain weight phenotypes across haplotypes and heterogeneity pseudo-genotypes from the two identified variants. The dots represent mean values and the error bars show standard error.

**Supplemental Figure 5.**
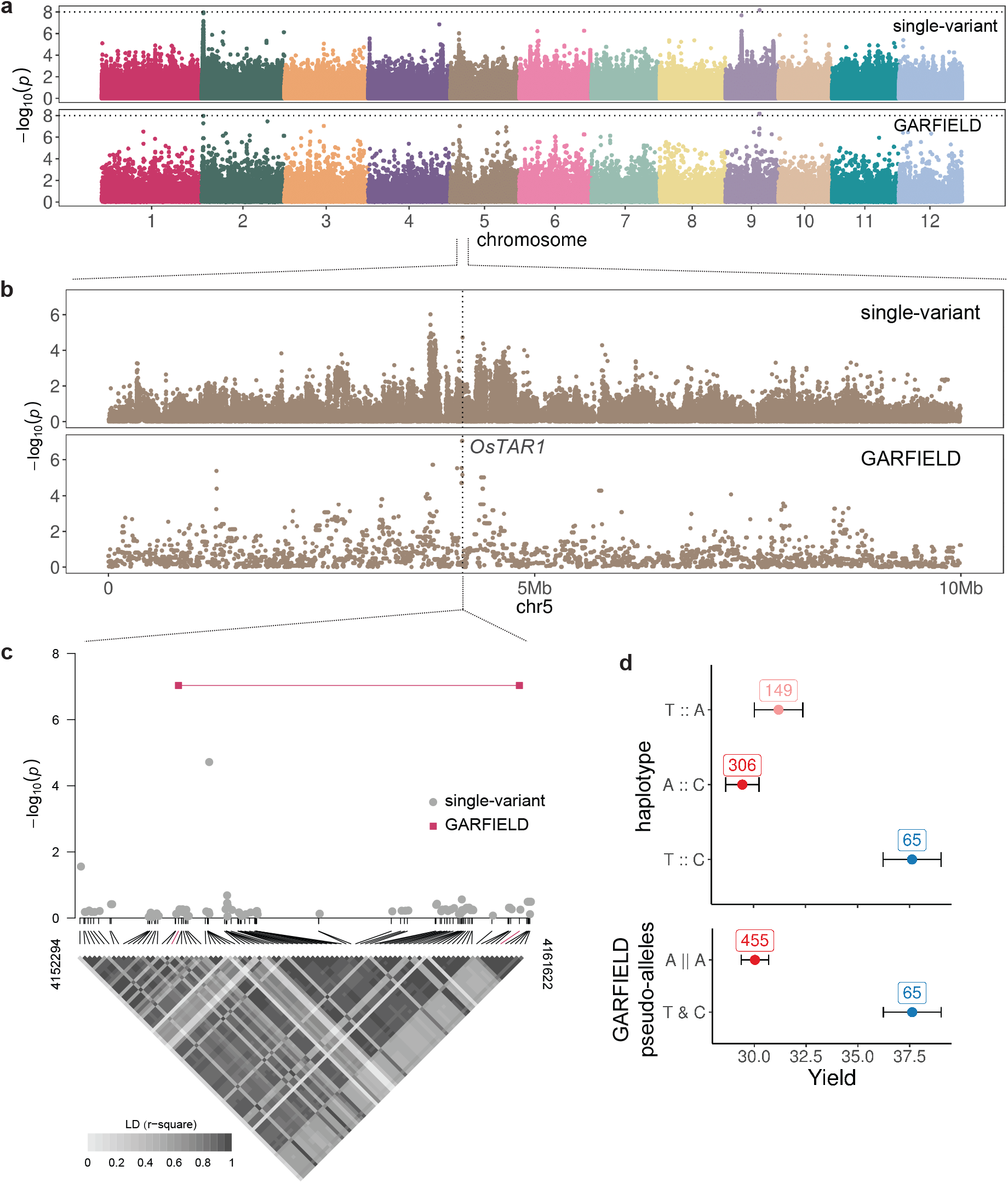
Allelic heterogeneity GWAS of rice yield identifies *OsTAR1*. **(a)** Comparison of single-locus GWAS and GARFIELD. **(b-c)** Zoom-in on *OsTAR1*. **(d)** The distribution of phenotypes across haplotypes and heterogeneity pseudo-genotypes resulting from the two identified variants. The A::A haplotype was excluded due to its presence in only 2 individuals. Dots show the mean, while error bars indicate standard error.

**Supplemental Figure 6.**
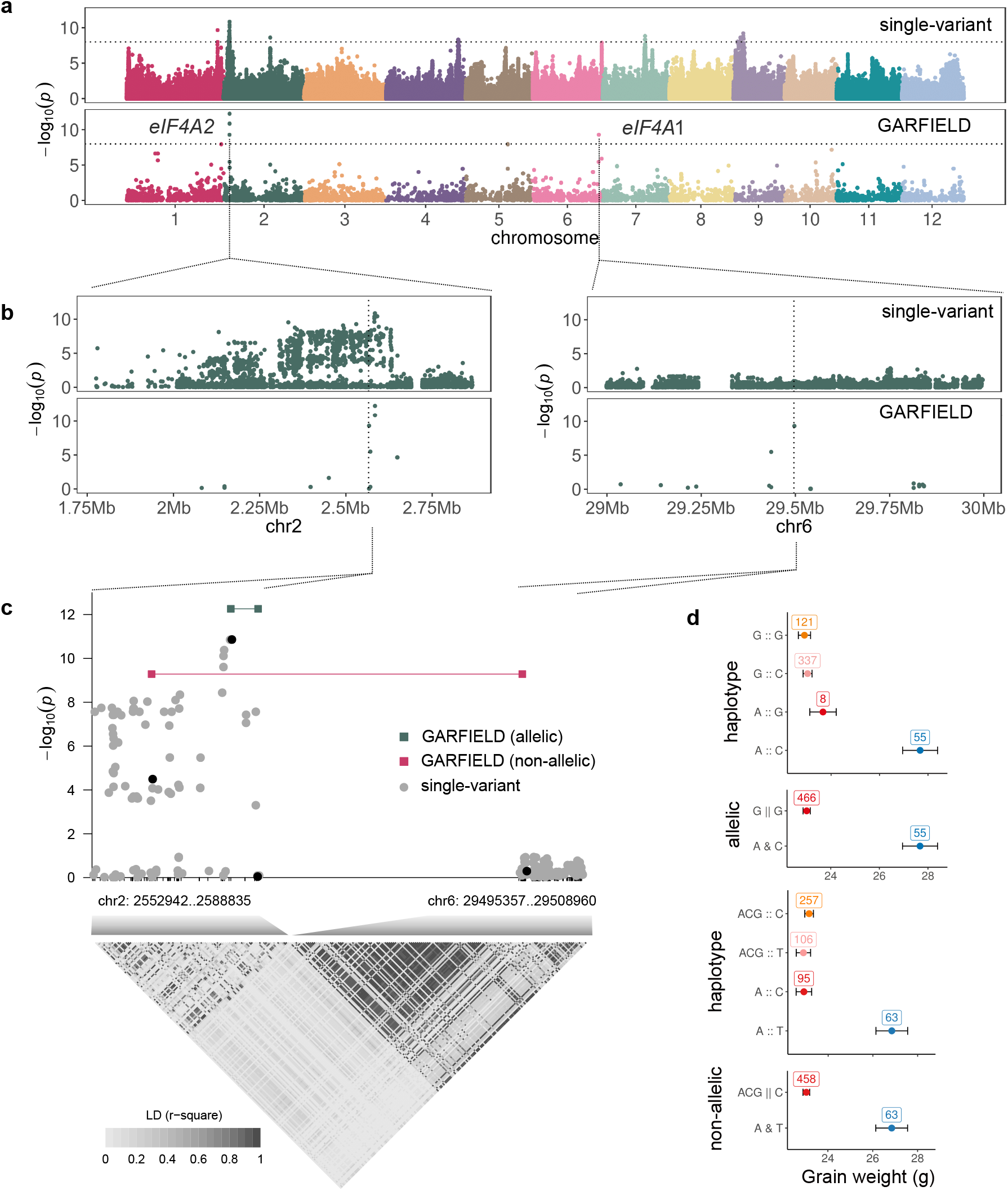
Allelic heterogeneity GWAS of rice grain weight identifies a novel allele of *OseIF4A2* locus and an interaction with its paralog, *OseIF4A1*. **(a)** Single-locus GWAS for rice grain weight compared to paralog-based non-allelic heterogeneity analysis. The former identifies many associations in a large region surrounding *OseIF4A2*, but nothing significant at the paralog *OseIF4A1*. **(b)** Heterogeneity analysis reveals a potential new allele at *OseIF4A2*. **(c)** Paralog-based heterogeneity analysis suggests an interaction between the two loci **(d)**. Summary of the single-locus and two-locus associations. **(e)** Variation in grain weight across haplotypes and heterogeneity pseudo-genotypes resulting from the interaction between the paralogs. Dots and error bars indicated the mean and standard error, respectively; with the sample size shown above each haplotype/allele.

**Supplemental Figure 7.**
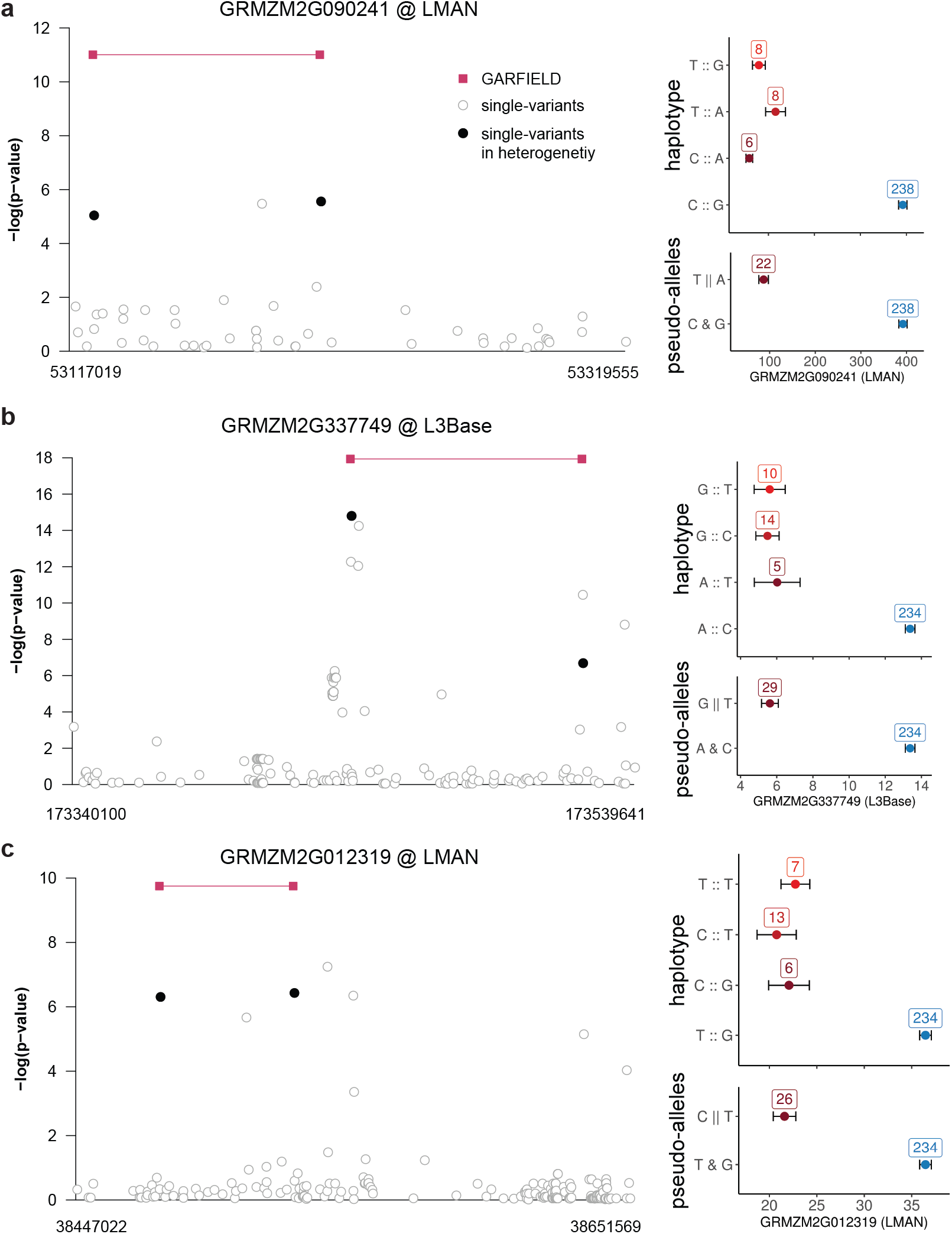
Examples of allelic heterogeneity eQTL in maize data. Allelic heterogeneity analysis revealed novel eQTL **(a)**, novel independent alleles **(b)**, and variant sets distinct from peaks detected through single-variant analysis **(c)**. The corresponding expression quantification across haplotypes and heterogeneity pseudo-genotypes is shown on the right. Dots show means and error bars show standard errors.

**Supplemental Figure 8.**
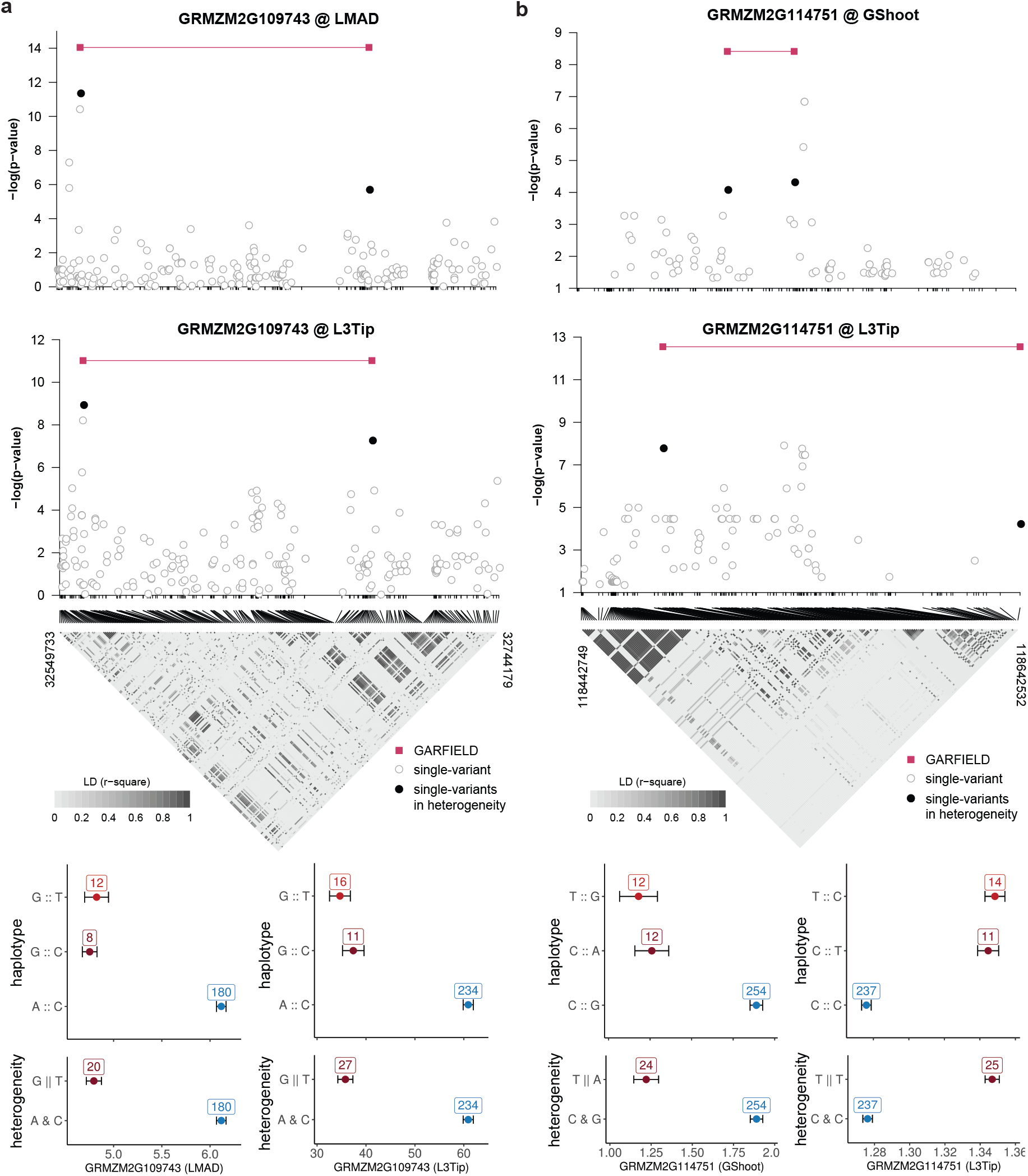
Instances of heterogeneity eQTLs for the same gene in different tissues. The sets of heterogeneity variants were either identical **(a)** or different **(b)** across various tissues. The corresponding expression quantification across haplotypes and heterogeneity pseudo-genotypes for the same gene in different tissues is illustrated in the lower panel. The dots and error bars indicate the mean and standard error, respectively.

**Supplemental Figure 9.**
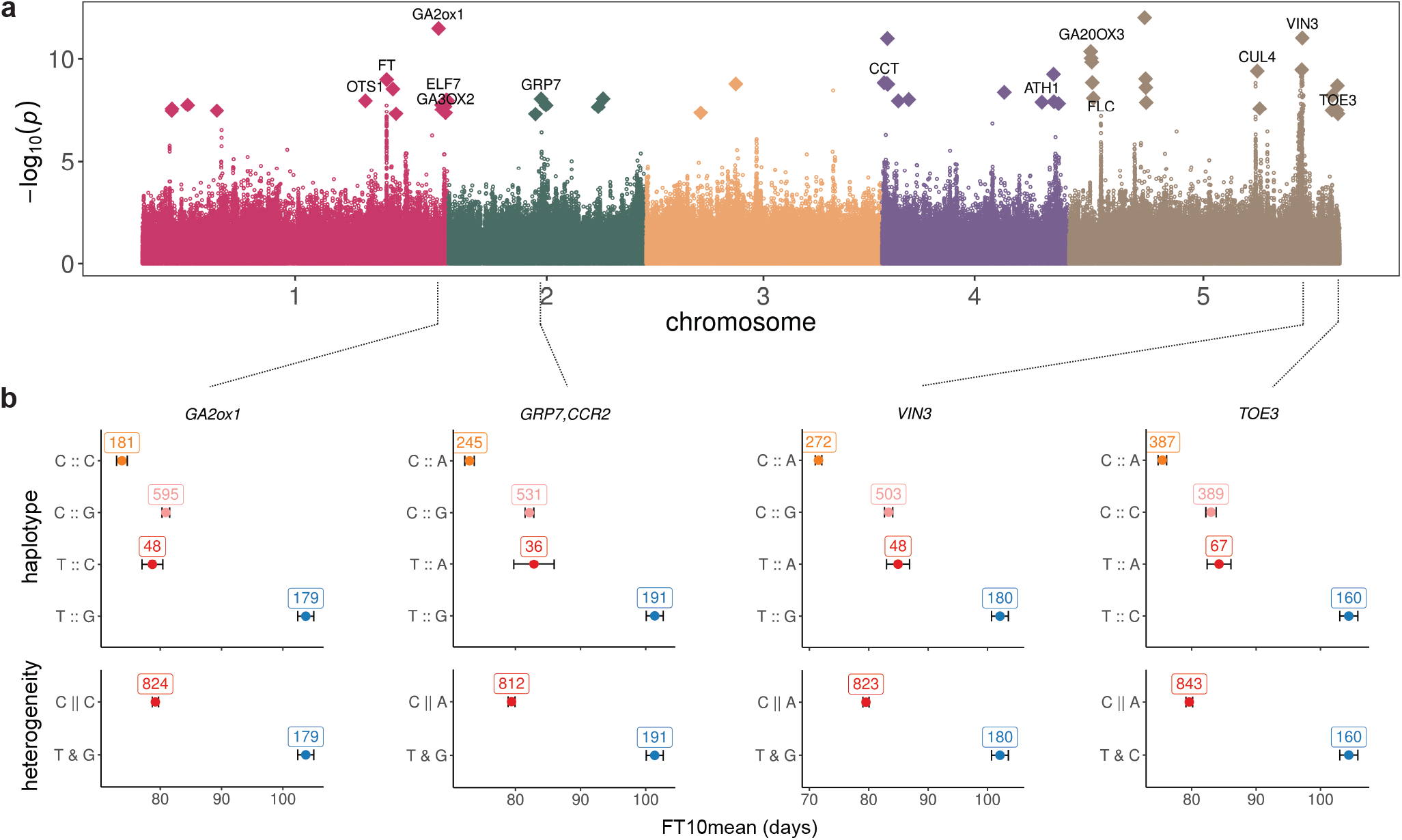
Identification of flowering time genes by non-allelic heterogeneity analysis on gene pairing with *FLC*. **(a)** GWAS for *A. thaliana* flowering time at 10^*°*^C^34^, with non-allelic heterogeneity results for interaction with *FLC* superimposed (as diamonds; only results with joint significance exceeding that of *FLC* alone are shown. **(b)** Flowering time variation across haplotypes and heterogeneity pseudo-genotypes resulting from the variant combinations between gene pairs. The four genes are indicated because their paired variants were found within their gene bodies, whereas the other candidate genes shown in (a) were positioned near the identified paired variants with *FLC*. Dots show means and error bars show standard errors.

**Supplemental Figure 10.**
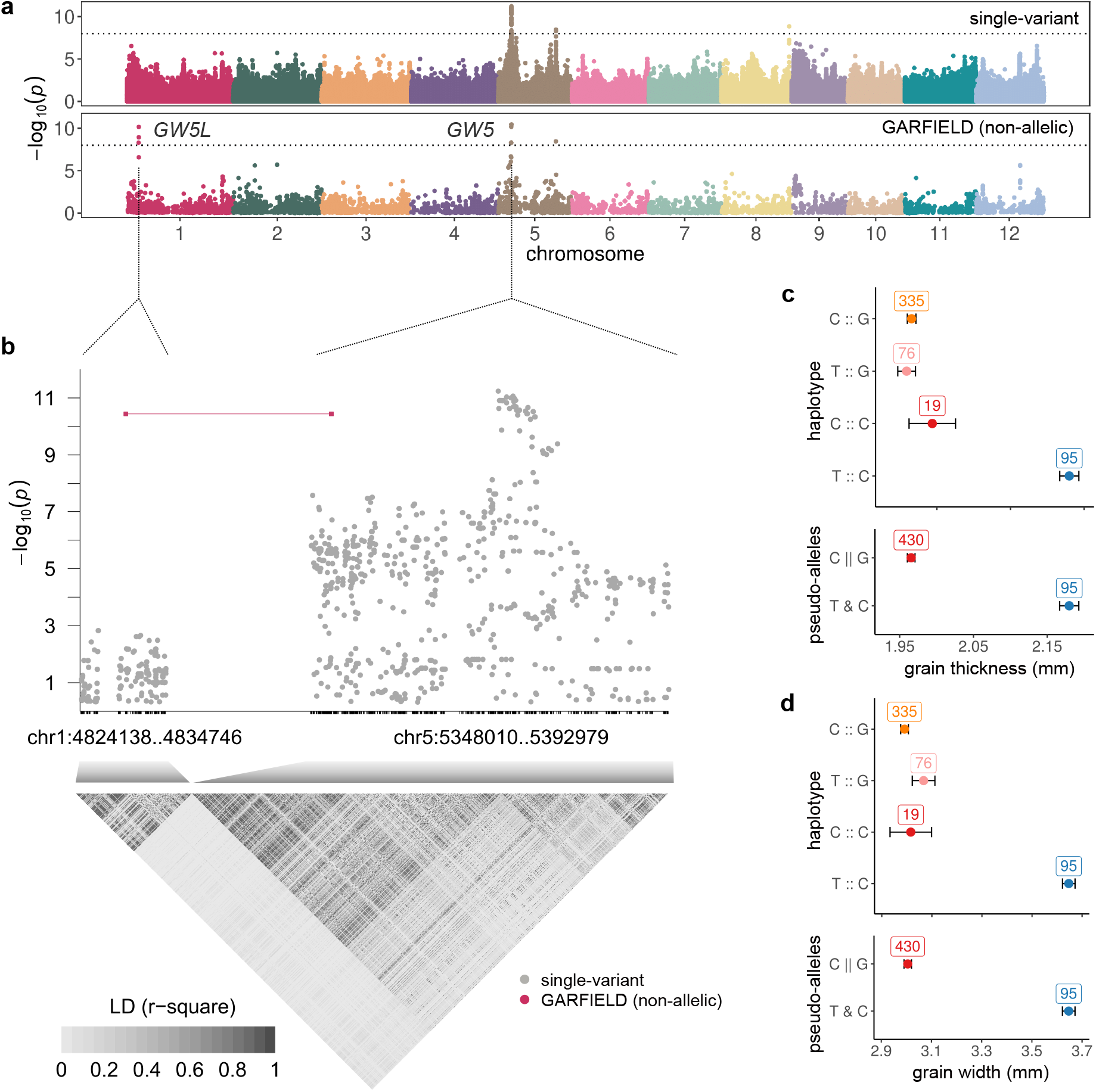
Identification of rice *GW5* paralogs associated with grain shape. **(a)** Mapping results for rice grain thickness using the single-variant-based and paralog-based heterogeneity methods. The regional plot **(b)** highlights the *GW5* paralogs revealed by the heterogeneity analysis. Grain thickness **(c)** and grain width variation **(d)** across haplotypes and heterogeneity pseudo-genotypes based on the combinations of paralog variants. Dots show means and error bars show standard errors.

**Supplemental Figure 11.**
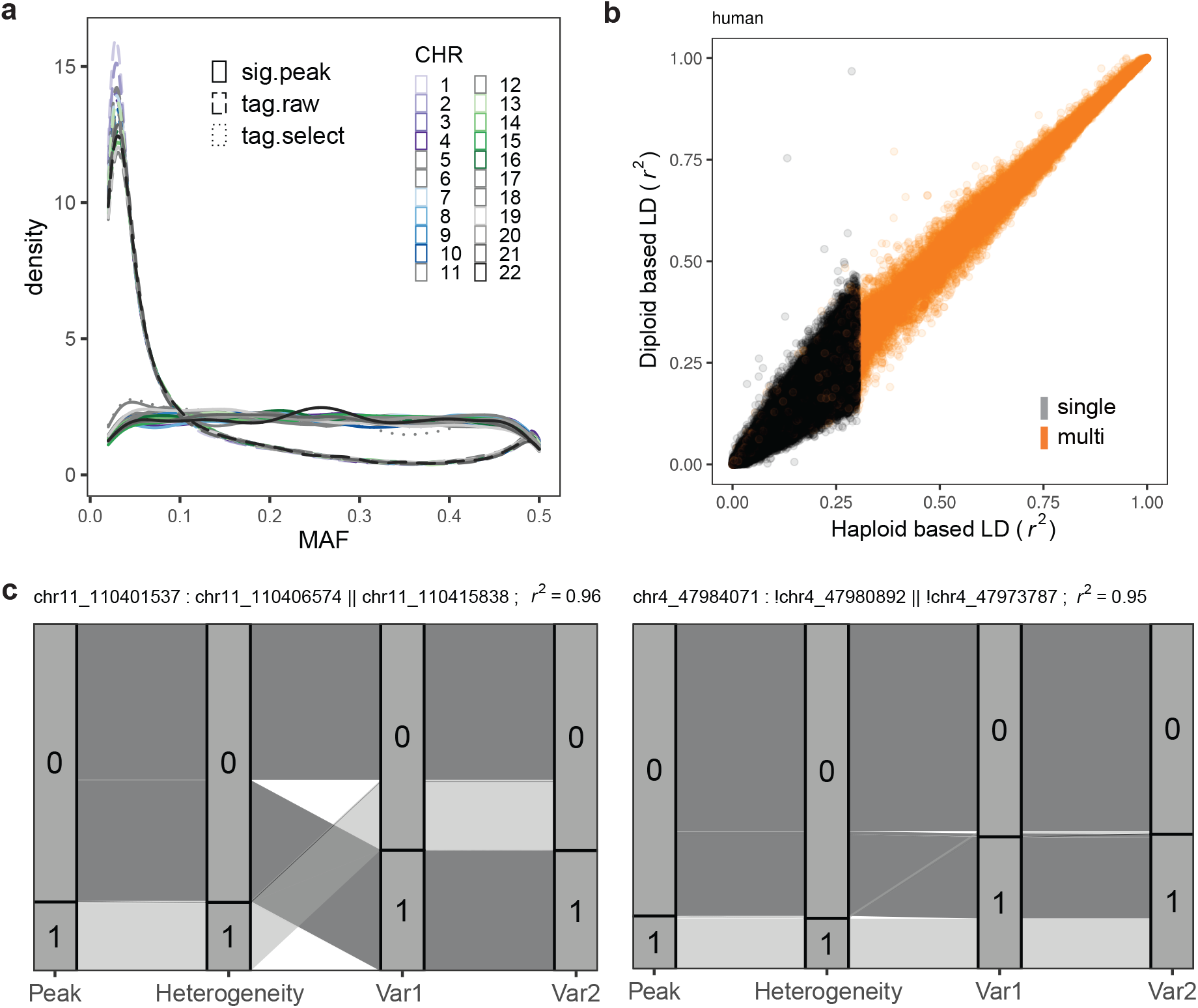
Technical issues in the analysis of human synthetic associations. **(a)** Distribution of minor allele frequency (MAF) for GWAS peaks, all tag variants selected only by LD pruning (tag.raw), and control tags selected to further match the MAF profiles of the GWAS peaks (tag.select). Colors denote chromosomes. **(b)** Correlation (*r*^2^) between GWAS peak variants (and GARFIELD pseudo-genotypes) generated from phased haplotypes (x-axis) and diploid individuals (y-axis). **(c)** Challenging scenarios were excluded from the analysis where different alleles of heterogeneity pseudo-genotypes exist in the same haplotype of the identified variant pair (Var1 and Var2). For example, the first case includes both 0 and 1 heterogeneity alleles in the 0:0 haplotype of Var1:Var2, while the second case shows both 0 and 1 heterogeneity alleles in the 1:1 haplotype of Var1:Var2.

**Supplemental Figure 12.**
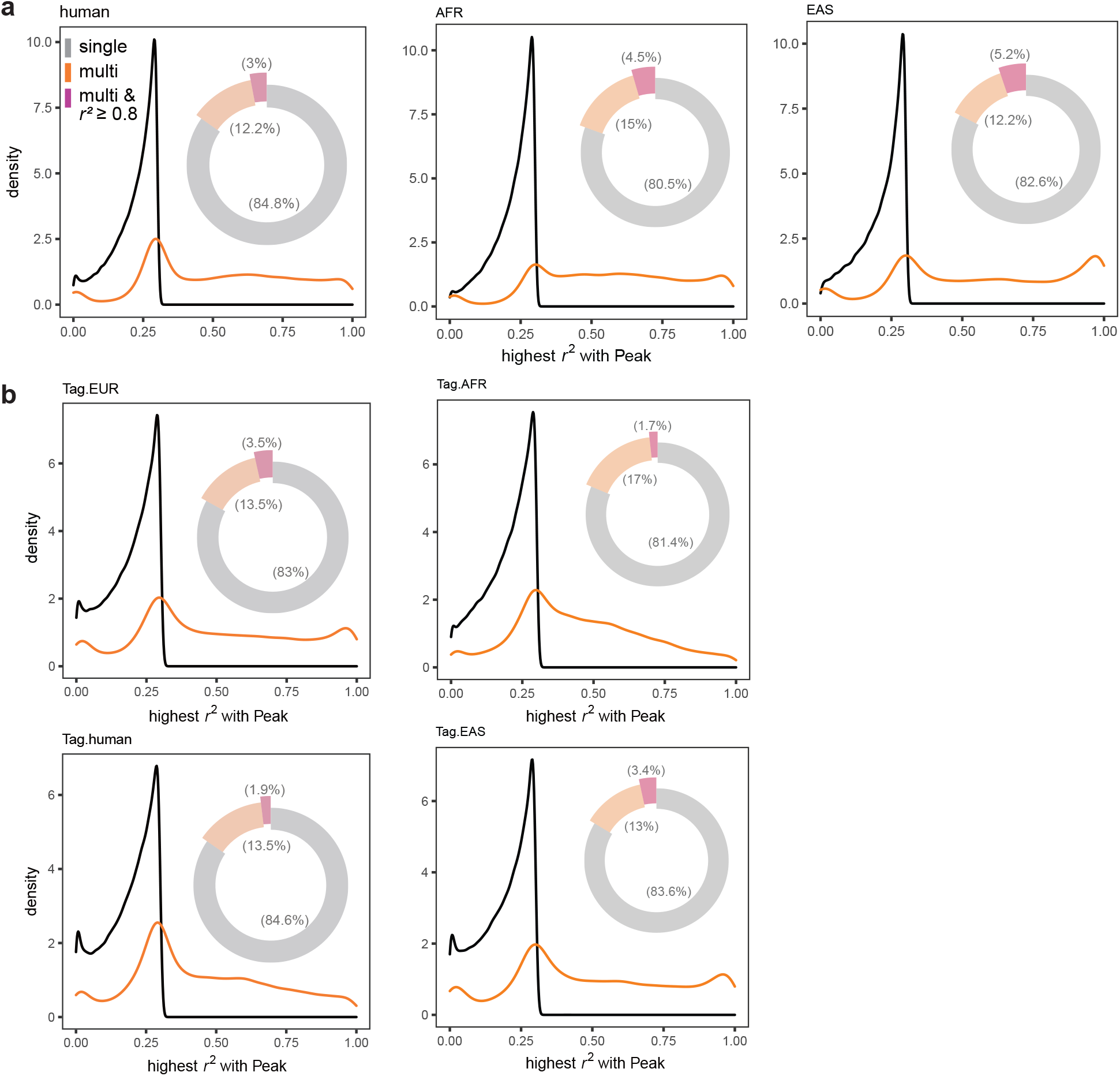
Identifying potential synthetic associations in human GWAS. **(a)** Same as Fig. 4b, but for the whole population (“human”) and the AFR and EAS subpopulations. **(b)** Same as Fig. 4b, but for randomly selected control “tag-variants” with matching MAF in the various populations.

**Supplemental Figure 13.**
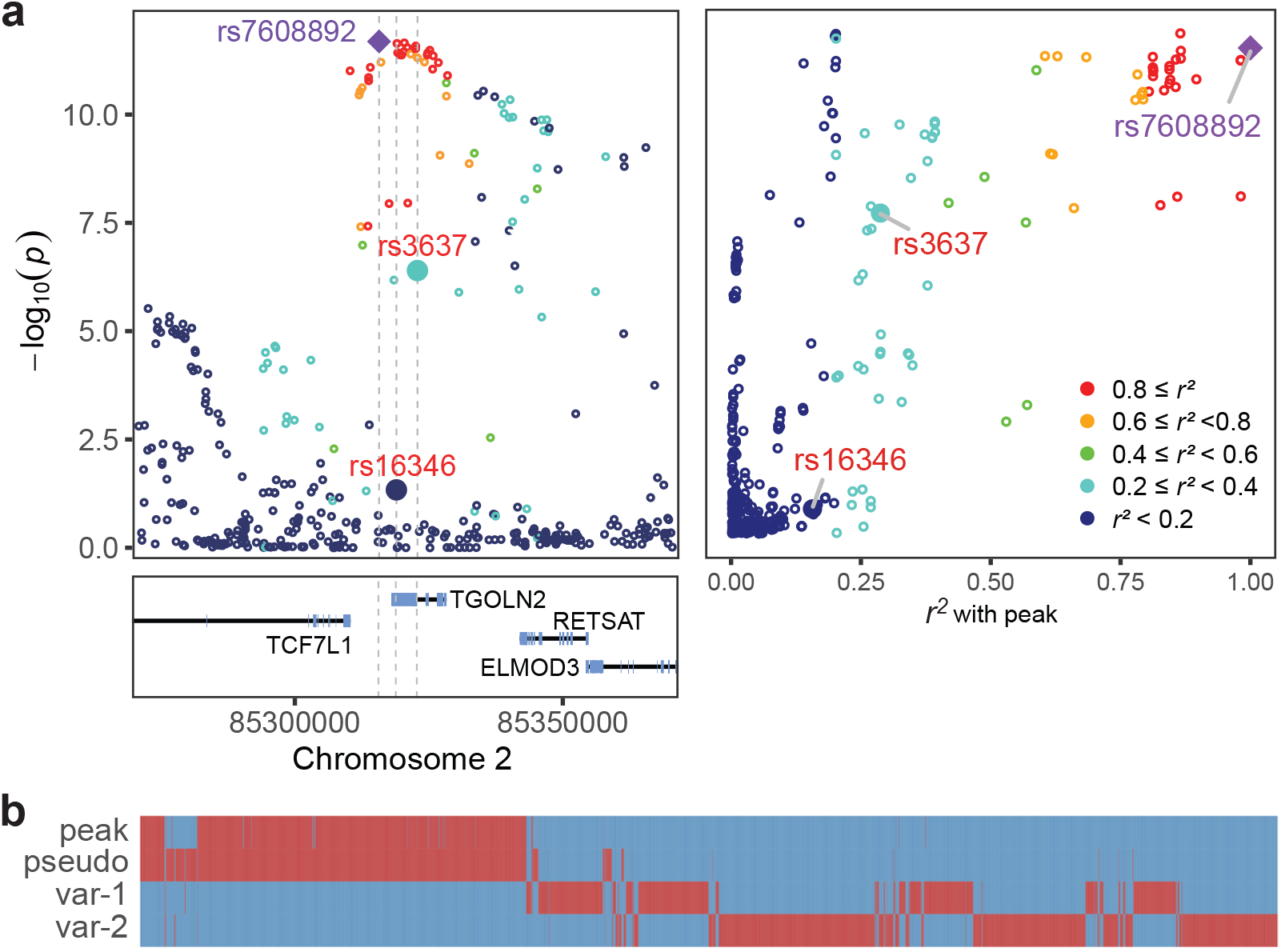
Potential SA in human GWAS at *TGOLN2* locus. **(a)** Potential SA of the *TGOLN2* locus on high-density lipoprotein cholesterol levels. Purple and red text indicate the GWAS peak and the underlying heterogeneity variants. Dot colors measure the LD level to the peak. **(b)** Detailed haplotypes within the *TGOLN2* region, alleles of the peak, heterogeneity pseudo-genotype, and the two variants consisting of the heterogeneity.

**Supplemental Figure 14.**
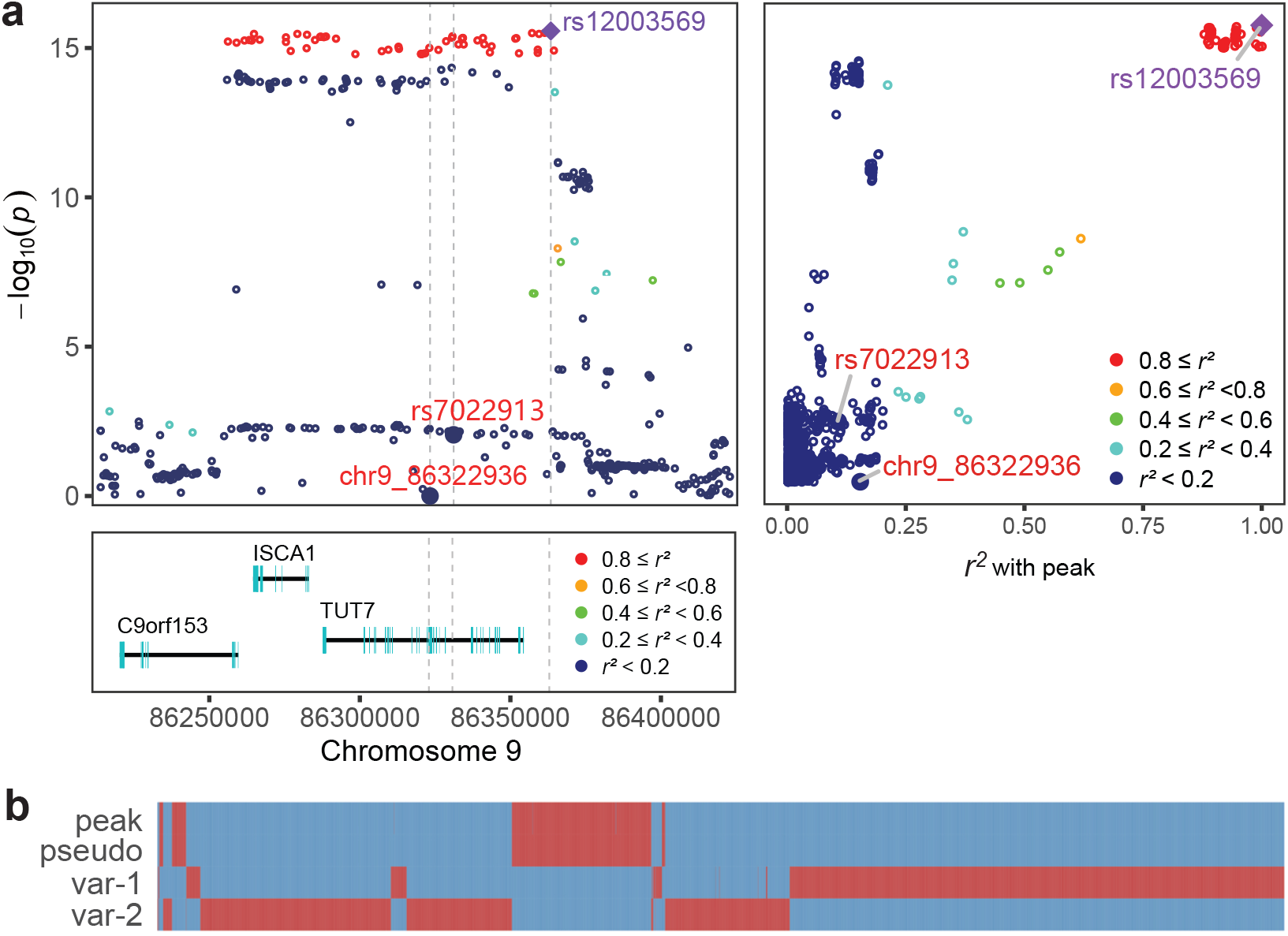
Potential SA at *TUT7* locus. **(a)** Potential SA of *TUT7* associated with cortical surface area. A disruptive inframe deletion and an intron SNP within this gene, when combined, exhibited a high correlation (LD *r*^*2*^ = 0.995) with the downstream intergenic peak. This likelihood was increased by the fact that the gene (RNU2-36P) downstream of the intergenic peak showed no expression in all measured GTEx samples, including various brain tissues. **(b)** Detailed haplotypes within the *TUT7* region, and allelic segregation of related genotypes.

**Supplemental Figure 15.**
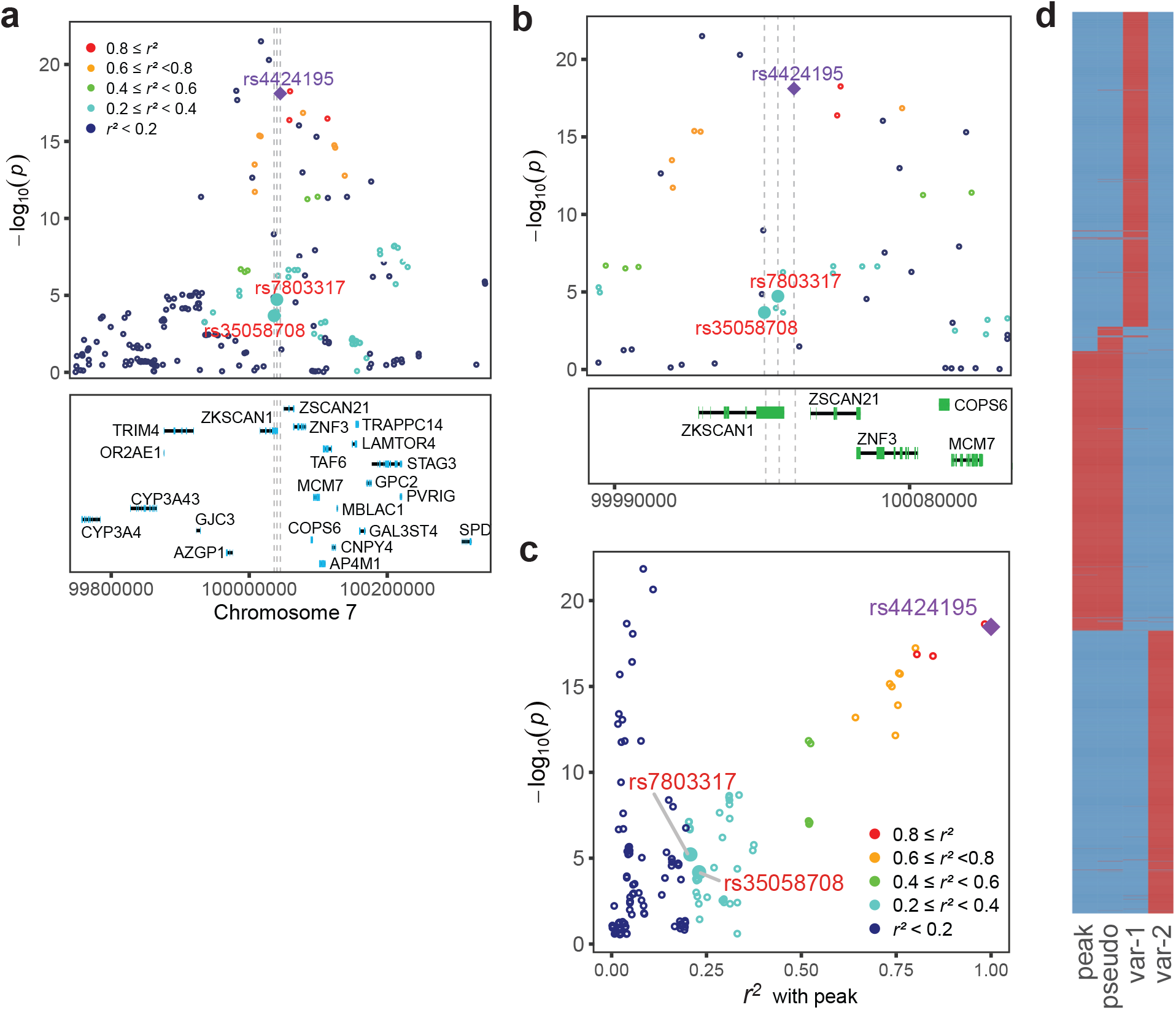
Potential SA of *ZKSCAN1* on body height. Potential SA of *ZKSCAN1* on body height, association in a larger **(a)** and more specific **(b)** region. **(c)** The relationship between p-values and LD to the peak within the *ZKSCAN1* region. **(d)** Detailed haplotypes of the *ZKSCAN1* region, and alleles of relevant genotypes.

**Supplemental Figure 16.**
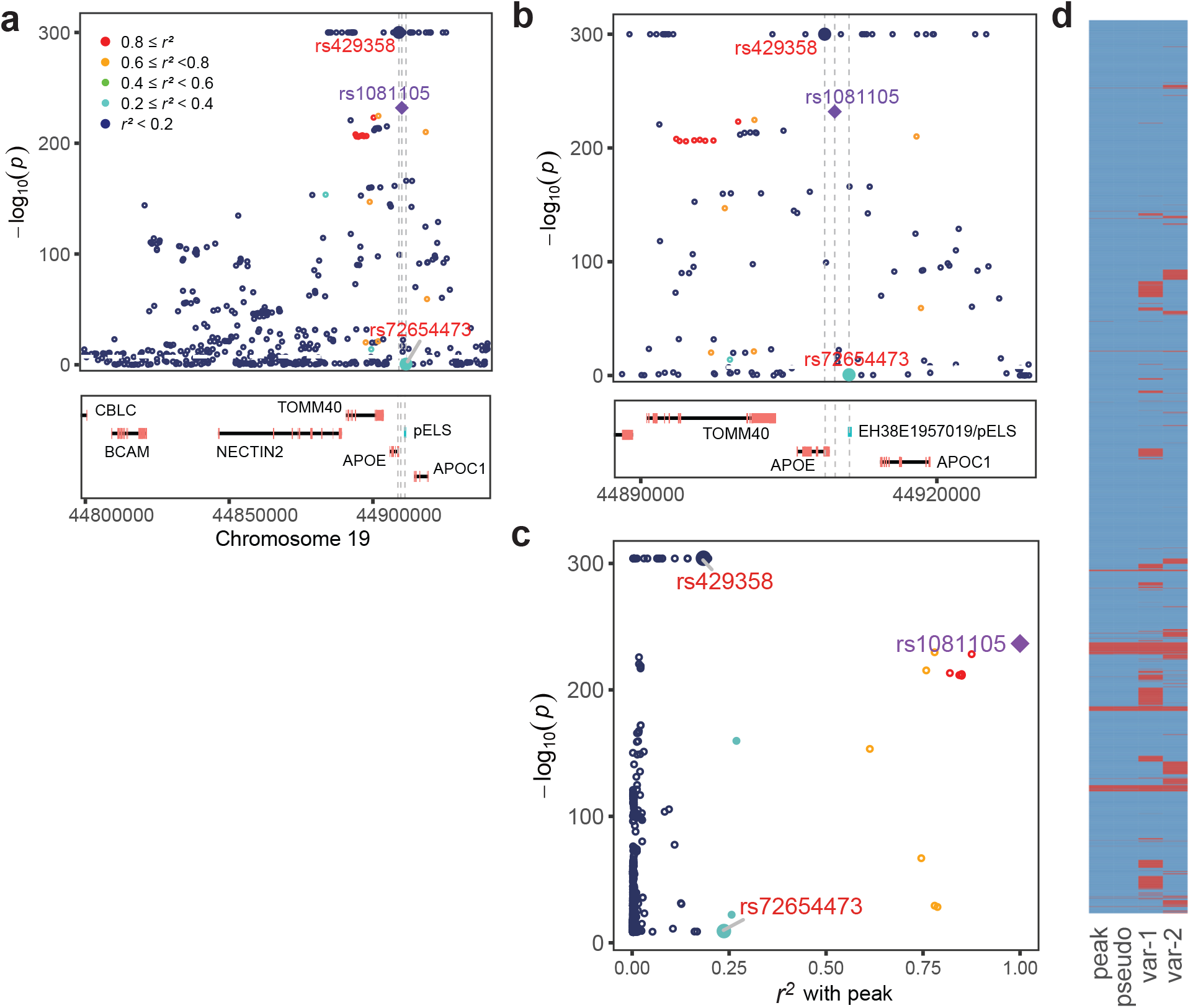
Potential misleading findings discovered at *APOE* locus to Alzheimer’s disease. Potential SA of the *APOE* locus for Alzheimer’s disease, a larger **(a)** and more specific **(b)** region. Note that one of the heterogeneity variants identified corresponds to the missense variant within *APOE* while the other resides in a potential cis-regulatory element (proximal enhancer-like signature, E1957019). The missense variant shows an extreme association with Alzheimer’s itself (*P* = 0 in the summary statistics and has been adjusted to *P* = 1×10^−300^ in the plot), but this variant, together with the others that displayed *P* = 0, was excluded in the corresponding studies. **(c)** The relationship between p-values and LD to the peak within the region. **(d)** Detailed haplotypes within the *APOE* region, and allelic segregation of related genotypes.

**Supplemental Figure 17.**
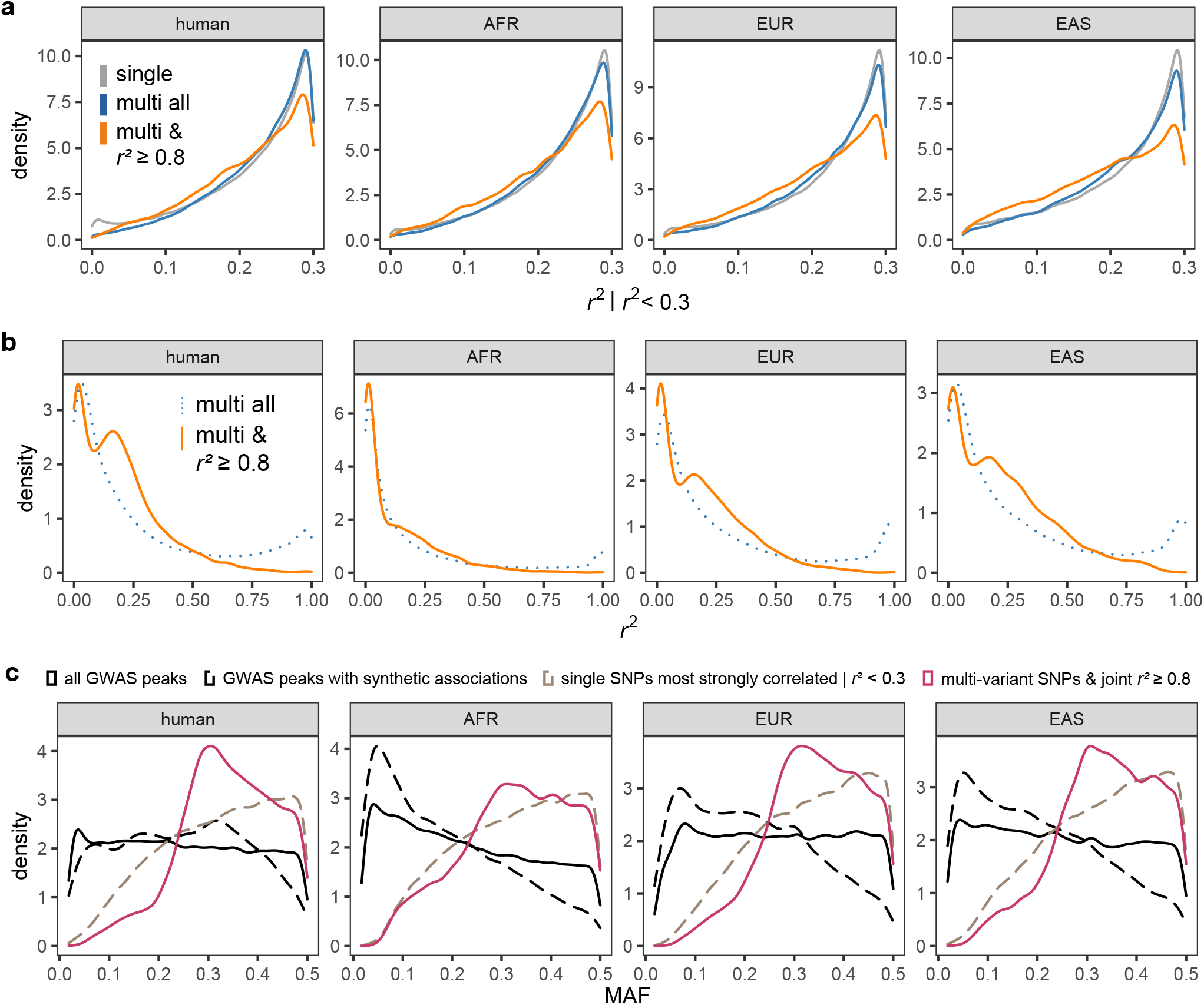
Additional features of potential synthetic associations in human GWAS. **(a)** The distribution of *r*^2^ between peak GWAS SNP and various sets of SNPs within 100 kb: most strongly correlate single variant, member of multi-variant sets, and multi-variants with joint *r*^2^≥0.8. Note that only SNPs with *r*^2^ *<* 0.3 are included, so the distribution is conditional on this. Plots show the three largest subpopulations and the full samples (“human”). **(b)** The distribution of *r*^2^ between multi-variant pseudo-genotypes and peak GWAS SNPs. Only cases with two identified variants are shown. **(c)** MAF distribution for different variant categories.

## Notes

### Competing Interest Statement

The authors have declared no competing interest.

## References

1. Uffelmann, E. et al. Genome-wide association studies. Nature Reviews Methods Primers 1, 1–21 (2021).

2. Tam, V. et al. Benefits and limitations of genome-wide association studies. Nat. Rev. Genet. 20, 467–484 (2019).

3. Woodward, A. A., Urbanowicz, R. J., Naj, A. C. & Moore, J. H. Genetic heterogeneity: Challenges, impacts, and methods through an associative lens. Genet. Epidemiol. 46, 555–571 (2022).

4. Dickson, S. P., Wang, K., Krantz, I., Hakonarson, H. & Gold-stein, D. B. Rare variants create synthetic genome-wide associations. PLoS Biol. 8, e1000294 (2010).

5. Platt, A., Vilhjálmsson, B. J. & Nordborg, M. Conditions under which genome-wide association studies will be positively misleading. Genetics 186, 1045–1052 (2010).

6. Lin, Z. et al. Parallel domestication of the Shattering1 genes in cereals. Nat. Genet. 44, 720–724 (2012).

7. Atwell, S. et al. Genome-wide association study of 107 pheno-types in Arabidopsis thaliana inbred lines. Nature 465, 627–631 (2010).

8. Kerdaffrec, E. et al. Multiple alleles at a single locus control seed dormancy in Swedish Arabidopsis. Elife 5, e22502 (2016).

9. Sasaki, E., Köcher, T., Filiault, D. L. & Nordborg, M. Revisiting a GWAS peak in Arabidopsis thaliana reveals possible confounding by genetic heterogeneity. Heredity 127, 245–252 (2021).

10. Huang, X. et al. Genome-wide association studies of 14 agronomic traits in rice landraces. Nat. Genet. 42, 961–967 (2010).

11. Barrett, J. C. et al. Genome-wide association defines more than 30 distinct susceptibility loci for Crohn’s disease. Nat. Genet. 40, 955–962 (2008).

12. Wray, N. R., Purcell, S. M. & Visscher, P. M. Synthetic associations created by rare variants do not explain most GWAS results. PLoS Biol. 9, e1000579 (2011).

13. Anderson, C. A., Soranzo, N., Zeggini, E. & Barrett, J. C. Synthetic associations are unlikely to account for many common disease genome-wide association signals. PLoS Biol. 9, e1000580 (2011).

14. Goldstein, D. B. The importance of synthetic associations will only be resolved empirically. PLoS Biol. 9, e1001008 (2011).

15. Sollis, E. et al. The NHGRI-EBI GWAS catalog: knowledge-base and deposition resource. Nucleic Acids Res. 51, D977– D985 (2023).

16. Abell, N. S. et al. Multiple causal variants underlie genetic associations in humans. Science 375, 1247–1254 (2022).

17. Breiman, L. Random forests. Mach. Learn. 45, 5–32 (2001).

18. Hormozdiari, F. et al. Widespread allelic heterogeneity in complex traits. Am. J. Hum. Genet. 100, 789–802 (2017).

19. Lopez-Arboleda, W. A., Reinert, S., Nordborg, M. & Korte, A. Global genetic heterogeneity in adaptive traits. Mol. Biol. Evol. 38, 4822–4831 (2021).

20. Hormozdiari, F., Jung, J., Eskin, E. & J Joo, J. W. MARS: leveraging allelic heterogeneity to increase power of association testing. Genome Biol. 22, 128 (2021).

21. Lee, S., Wu, M. C. & Lin, X. Optimal tests for rare variant effects in sequencing association studies. Biostatistics 13, 762–775 (2012).

22. Li, Z. et al. A framework for detecting noncoding rare-variant associations of large-scale whole-genome sequencing studies. Nat. Methods 19, 1599–1611 (2022).

23. Lunetta, K. L., Hayward, L. B., Segal, J. & Van Eerdewegh, P. Screening large-scale association study data: exploiting interactions using random forests. BMC Genet. 5, 32 (2004).

24. Bureau, A. et al. Identifying SNPs predictive of phenotype using random forests. Genet. Epidemiol. 28, 171–182 (2005).

25. Li, H. et al. Genome-wide association study dissects the genetic architecture of oil biosynthesis in maize kernels. Nat. Genet. 45, 43–50 (2013).

26. Liu, H. et al. Distant eQTLs and non-coding sequences play critical roles in regulating gene expression and quantitative trait variation in maize. Mol. Plant 10, 414–426 (2017).

27. Baud, S., Wuillème, S., To, A., Rochat, C. & Lepiniec, L. Role of WRINKLED1 in the transcriptional regulation of glycolytic and fatty acid biosynthetic genes in Arabidopsis. Plant J. 60, 933–947 (2009).

28. Pouvreau, B. et al. Duplicate maize wrinkled1 transcription factors activate target genes involved in seed oil biosynthesis. Plant Physiol. 156, 674–686 (2011).

29. Asano, T. et al. Rice SPK, a calmodulin-like domain protein kinase, is required for storage product accumulation during seed development: phosphorylation of sucrose synthase is a possible factor. Plant Cell 14, 619–628 (2002).

30. Xu, X. et al. OsYUC11-mediated auxin biosynthesis is essential for endosperm development of rice. Plant Physiol. 185, 934–950 (2020).

31. Kremling, K. A. G. et al. Dysregulation of expression correlates with rare-allele burden and fitness loss in maize. Nature 555, 520–523 (2018).

32. Rodgers-Melnick, E., Vera, D. L., Bass, H. W. & Buckler, E. S. Open chromatin reveals the functional maize genome. Proc. Natl. Acad. Sci. U. S. A. 113, E3177–84 (2016).

33. Bouch é, F., Lobet, G., Tocquin, P. & Périlleux, C. FLOR-ID: an interactive database of flowering-time gene networks in arabidopsis thaliana. Nucleic Acids Res. 44, D1167–D1171 (2015).

34. 1001 Genomes Consortium. 1,135 genomes reveal the global pattern of polymorphism in arabidopsis thaliana. Cell 166, 481–491 (2016).

35. Peng, J. Gene redundancy and gene compensation: An updated view. J. Genet. Genomics 46, 329–333 (2019).

36. Parrish, P. C. R. et al. Discovery of synthetic lethal and tumor suppressor paralog pairs in the human genome. Cell Rep. 36, 109597 (2021).

37. Gonatopoulos-Pournatzis, T. et al. Genetic interaction mapping and exon-resolution functional genomics with a hybrid Cas9-Cas12a platform. Nat. Biotechnol. 38, 638–648 (2020).

38. Weng, J. et al. Isolation and initial characterization of GW5, a major QTL associated with rice grain width and weight. Cell Res. 18, 1199–1209 (2008).

39. Shomura, A. et al. Deletion in a gene associated with grain size increased yields during rice domestication. Nat. Genet. 40, 1023–1028 (2008).

40. Duan, P. et al. Natural variation in the promoter of GSE5 contributes to grain size diversity in rice. Mol. Plant 10, 685–694 (2017).

41. Tian, P. et al. GW5-Like, a homolog of GW5, negatively regulates grain width, weight and salt resistance in rice. J. Integr. Plant Biol. 61, 1171–1185 (2019).

42. Byrska-Bishop, M. et al. High-coverage whole-genome sequencing of the expanded 1000 genomes project cohort including 602 trios. Cell 185, 3426–3440.e19 (2022).

43. Graham, S. E. et al. The power of genetic diversity in genome-wide association studies of lipids. Nature 600, 675–679 (2021).

44. Saunders, G. R. B. et al. Genetic diversity fuels gene discovery for tobacco and alcohol use. Nature 612, 720–724 (2022).

45. Matsui, A. et al. BTBD3 controls dendrite orientation toward active axons in mammalian neocortex. Science 342, 1114–1118 (2013).

46. Thompson, S. L. et al. Btbd3 expression regulates compulsive-like and exploratory behaviors in mice. Transl. Psychiatry 9, 222 (2019).

47. Gillies, P. J. et al. Regulation of inflammatory and lipid metabolism genes by eicosapentaenoic acid-rich oil. J. Lipid Res. 53, 1679–1689 (2012).

48. Davydov, E. V. et al. Identifying a high fraction of the human genome to be under selective constraint using GERP++. PLoS Comput. Biol. 6, e1001025 (2010).

49. Pollard, K. S., Hubisz, M. J., Rosenbloom, K. R. & Siepel, A. Detection of nonneutral substitution rates on mammalian phylogenies. Genome Res. 20, 110–121 (2010).

50. Hill, W. G., Goddard, M. E. & Visscher, P. M. Data and theory point to mainly additive genetic variance for complex traits. PLoS Genet. 4, e1000008 (2008).

51. Stephan, J., Stegle, O. & Beyer, A. A random forest approach to capture genetic effects in the presence of population structure. Nat. Commun. 6, 1–10 (2015).

52. Wright, M. N. & Ziegler, A. ranger: A fast implementation of random forests for high dimensional data in C++ and R. J. Stat. Softw. 77, 1–17 (2017).

53. Schwender, H. & Ickstadt, K. Identification of SNP interactions using logic regression. Biostatistics 9, 187–198 (2008).

54. Kang, H. M. et al. Variance component model to account for sample structure in genome-wide association studies. Nat. Genet. 42, 348–354 (2010).

55. Chang, C. C. et al. Second-generation PLINK: rising to the challenge of larger and richer datasets. Gigascience 4, s13742.– 015–0047–8 (2015).

56. Li, M.-X., Yeung, J. M. Y., Cherny, S. S. & Sham, P. C. Evaluating the effective numbers of independent tests and significant p-value thresholds in commercial genotyping arrays and public imputation reference datasets. Hum. Genet. 131, 747–756 (2012).

57. Yang, J., Lee, S. H., Goddard, M. E. & Visscher, P. M. GCTA: a tool for genome-wide complex trait analysis. Am. J. Hum. Genet. 88, 76–82 (2011).

58. Nicholls, H. L. et al. Reaching the End-Game for GWAS: Machine learning approaches for the prioritization of complex disease loci. Front. Genet. 11, 350 (2020).

59. Cordell, H. J. Detecting gene-gene interactions that underlie human diseases. Nat. Rev. Genet. 10, 392–404 (2009).

60. Liu, H. et al. MODEM: multi-omics data envelopment and mining in maize. Database 2016 (2016).

61. Browning, B. L., Zhou, Y. & Browning, S. R. A One-Penny imputed genome from Next-Generation reference panels. Am. J. Hum. Genet. 103, 338–348 (2018).

62. Zhao, H. et al. RiceVarMap: a comprehensive database of rice genomic variations. Nucleic Acids Res. 43, D1018–22 (2015).

63. Ramstein, G. P. & Buckler, E. S. Prediction of evolutionary constraint by genomic annotations improves functional prioritization of genomic variants in maize. Genome Biol. 23, 183 (2022).

